# Improved state change estimation in dynamic functional connectivity using hidden semi-Markov models

**DOI:** 10.1101/519868

**Authors:** Heather Shappell, Brian S. Caffo, James J. Pekar, Martin A. Lindquist

## Abstract

The study of functional brain networks has grown rapidly over the past decade. While most functional connectivity (FC) analyses estimate one static network structure for the entire length of the functional magnetic resonance imaging (fMRI) time series, recently there has been increased interest in studying time-varying changes in FC. Hidden Markov models (HMMs) have proven to be a useful modeling approach for discovering repeating graphs of interacting brain regions (brain states). However, a limitation lies in HMMs assuming that the sojourn time, the number of consecutive time points in a state, is geometrically distributed. This may encourage inaccurate estimation of the time spent in a state before switching to another state. We propose a hidden semi-Markov model (HSMM) approach for inferring time-varying brain networks from fMRI data, which explicitly models the sojourn distribution. Specifically, we propose using HSMMs to find each subject’s most probable series of network states and the graphs associated with each state, while properly estimating and modeling the sojourn distribution for each state. We perform a simulation study, as well as an analysis on both task-based fMRI data from an anxiety-inducing experiment and resting-state fMRI data from the Human Connectome Project. Our results demonstrate the importance of model choice when estimating sojourn times and reveal their potential for understanding healthy and diseased brain mechanisms.

## 1. Introduction

Interest in studying brain networks, where the brain is viewed as a system of interacting regions (nodes) that produce complex behaviors, has grown tremendously over the past decade. Networks have become a popular approach towards illustrating both the physiological connections (structural networks) and the coupling of dynamic brain activity (functional networks) linking different areas of the brain. It is within this paradigm shift that scientists have begun investigating how networks behave in healthy brains and how they are altered in neurological and psychiatric disorders [1].

The study of functional connectivity (FC), or the undirected association between two or more fMRI time series, has come to the forefront of research efforts in the field of neuroimaging. Studies have revealed information about the functional connections between different brain regions and local networks and have provided significant insights into the organization of functional communication in the brain. Brain networks can be created by performing a FC analysis, where the strength of the relationship between nodes is assessed using the functional magnetic resonance imaging (fMRI) time series associated with each node. The nodes can consist of individual voxels, [2, 3, 4], pre-specified regions of interest (ROIs) [5], or sets of regions estimated using independent component analysis (ICA) [6]. The relationship between nodes can be quantified using a variety of metrics, and we present the networks in this paper using both Pearson and partial correlation, as both are popular approaches.

To date, much of the brain connectivity analyses performed estimate one network state that does not change over time during a specific experimental scanning session. In other words, the network summarizes an aggregate state over time. However, recently there has been a growing interest in studying changes in connectivity over shorter time periods, i.e. on the second-to-minute scale [3, 7]. It is now generally believed that individuals transition in and out of different brain states during the course of the scanning session. Thus, there is a need to estimate the timing of the transitions between states, as well as the structure of the states themselves. Several approaches towards assessing time-varying connectivity have been suggested. These include using sliding window correlations [6], change point models [8, 9, 10], wavelet transform coherence [5], and time series models [11, 12].

There is a recent literature on Markov switching vector autoregressive (SVAR) models and dynamic connectivity. Samdin et al. [13] developed a framework in which the dynamic connectivity systems are characterized by distinct VAR processes and allowed to switch between brain states, where the state evolution and directed dependencies are described by a Markov process and the SVAR parameters. They demonstrate that their approach is able to detect change points and provide reliable estimates of effective connectivity. In addition, Ting et al. [14] propose a Markov-switching dynamic factor model, allowing for the dynamic network states to be driven by latent factors. A regime-switching SVAR factor process quantifies the dynamic effective connectivity. Lastly, Ombao et al. [15] present approaches to modeling dynamic connectivity using EEG data recorded across replicated trials in an experiment.

In recent years, Hidden Markov models (HMMs) have also proven to be a useful modeling approach towards assessing FC. In this context, it is assumed that the time series data at each brain region can be described via a series of a fixed number of hidden/unknown brain states. Each state is characterized by a multivariate Gaussian distribution, which is parameterized by the mean and covariance. Of particular interest is the unique connectivity structure of each state, encapsulated within the covariance matrix.

HMMs [16] are a natural modeling approach to take, given that we have an observable sequence from a system in which it is not unreasonable to believe that the hidden states may be governed by a Markov process. They have been widely studied in statistics [17] and have been applied extensively in many applications, such as in speech recognition and biological sequence analysis [18, 19]. In the context of dynamic FC, Eavani et al. [20] introduced a HMM framework to uncover hidden states in resting-state fMRI data. They simultaneously modeled the underlying co-varying regions of interest in each state as a set of sparse rank-one basis matrices, such that non-negative combinations of these basis matrices act as priors for each of the HMM covariance matrices. Vidaurre et al. [21] also utilized a HMM based analysis to discover a hierarchically organized temporal nature of resting-state networks. They showed that resting-state brain networks are organized into two metastates, where states within a metastate are highly likely to transition to other states within the same metastate. The authors provide evidence that the metastates are consistent across subjects, associated with behavior, and heritable. Moreover, Baker et al. [22] applied HMMs to band-limited amplitude envelopes of source reconstructed MEG data and identified brain states that match up to previously established resting-state networks and which fluctuate at time scales two orders of magnitude faster than previously shown. Warnick et al. [23] introduce a Bayesian approach to HMMs and dynamic functional connectivity, where they identify latent network states, but they do so in a manner where they assume the network/connectivity structures at each time point are related within one “super-graph”. They impose a sparsity inducing Markov random field prior on the presence of the edges in the super-graph.

A HMM approach to dynamic FC analysis is appealing for several reasons. First, the framework is able to handle large amounts of data to infer the network states, as well as the other model parameters such as the probabilities of transitioning between states. Moreover, these states and parameters can be estimated in a relatively computationally inexpensive manner. Second, while network states are estimated at the group level, individuality is respected in that information about when a state becomes active is subject-specific. Therefore, we are still able to obtain subject-specific estimates of an individual’s state sequence and the amount of time he/she spends in each state. Third, the method is likelihood based, leaving room for model selection techniques to be employed when deciding upon the number of states to be estimated, as well as the number of brain regions to be included in the analysis. Fourth, the model is able to capture quick changes in brain connectivity, unlike the sliding window based approach.

Unfortunately, with all of the aforementioned strengths, comes a limitation and potentially unrealistic assumption of the HMM approach, which is that the sojourn time, the number of consecutive time points spent in a specific state, is geometrically distributed. A property of the geometric distribution is that the probability decreases as the sojourn time increases. Therefore, more weight is placed on shorter consecutive time points in a given state. This may not be appropriate in the context of functional connectivity analysis, as it suggests subjects are switching states very often, as opposed to potentially spending longer periods of time in the same state. For example, in some task-based fMRI experiments, participants are expected to spend several minutes in a particular brain state. It also suggests that the sojourn time for all states follow a very similar pattern, which may not be the case in practice.

A possible solution to this issue is to explicitly model and estimate the state sojourn distributions via Hidden semi-Markov models (HSMMs) [24]. HSMMs differ in that the sojourn distributions are explicitly built into the likelihood function. One may specify the distributions to take a nonpara-metric form or a parametric form (such as a Poisson or Gamma distribution) and estimate the parameters directly. A more detailed comparison of HMMs vs. HSMMs can be found in Sections 2.4 and 2.5.

In this work, we propose using a HSMM approach to infer dynamic FC networks from fMRI data. We are particularly interested in modeling group level data in order to estimate several network states that are common across subjects in a particular study. Moreover, the parameter estimates we obtain from fitting the HSMM allow us to find each individual subject’s most probable series of network states along with the amount of time he/she is in each state. The estimated model also lends additional information, such as the probabilities of transitioning from one network state to another, the sojourn distribution of each state, and the initial state probabilities.

We first conduct a simulation study, revealing that while both the HMM and HSMM are able to identify quick state switches with similar accuracy, the HSMM is superior to the HMM with regards to estimating sojourn times with fewer switches. This suggests that the HSMM is a more flexible model and the better model choice when estimating sojourn times and subject state sequences for data where subjects aren’t expected to switch states at a rapid rate. We then demonstrate our approach on an anxiety-inducing experiment [25, 26], in which the algorithm was agnostic to alignment, and yet discovered the alignment near perfectly. These results suggest reliability of the method, which is quite encouraging for moving forward with analyzing resting-state data. Lastly, we present dynamic FC results using resting-state fMRI data from the Human Connectome Project (HCP), where we identify several states associated with different degrees of connectedness within and between sensory, motor, and higher order cognition regions. Moreover, we find that the sojourn distributions of several states are associated with scores on a sustained attention task, highlighting how informative they may be in understanding healthy and diseased brain mechanisms. Throughout the paper we make comparisons to the traditional HMM approach, illustrating that results can differ depending on which method is used.

## 2. Materials and Methods

### 2.1 Anxiety Inducing Experiment Data and Preprocessing

We fit our HMM and HSMM on 23 participants who performed an anxiety-inducing task while in the fMRI scanner [27]. The experiment, approved by the University of Michigan institutional review board, was conducted in accordance with the Declaration of Helsinki.

The experimental paradigm was an off-on-off design, with an anxiety-provoking task occurring between two resting periods. In particular, participants were informed prior to the scan that they were to be given 2 minutes to prepare a 7 minute speech and that the topic would be revealed to them during the scanning session. They were told that after the scan they would be asked to present the speech to a panel of expert judges, with there being “a small chance” that they would be randomly selected not to give the speech.

Each scan began with the participants viewing a fixation cross for 2 minutes (resting baseline). Next, an instruction slide was shown for 15 seconds that described the speech topic, i.e.”why you are a good friend”. After the 2 minutes allotted for the speech preparation, another instruction screen appeared for 15 seconds that informed participants that they would *not* have to give the speech. Lastly, the scan ended with an additional 2 minute period of resting baseline. Heart rate was monitored for the entire length of the scan.

Throughout the course of the experiment, a series of 215 functional images were acquired (TR = 2 s). A detailed description of the data acquisition and preprocessing can be found in previous work [27]. Time series of voxels were averaged across pre-specified ROIs. We used data consisting of 4 ROIs and heart rate. The 4 ROIs were chosen based on the fact that they showed a significant relationship to heart rate in an independent data set. The ROIs included the ventral medial prefrontal cortex (VMPFC), the anterior medial prefrontal cortex (mPFC), the striatum/pallidum, and the dorsal lateral prefrontal cortex (DLPFC)/inferior frontal junction (IFJ). The temporal resolution of the heart rate was 1 second compared to 2 seconds for fMRI data, so it was down-sampled by taking every other measurement.

### 2.2 HCP Data and Pre-processing

We also analyzed resting-state fMRI data from 820 subjects from the Human Connectome Project (HCP). The HCP provides the required ethics and consent needed for study and dissemination. Therefore, no further institutional review board approval is required. The 820 subjects all have complete resting-state fMRI data from the 900-subject public data release. The sample includes all healthy adults (22 - 35 years old, 453 females) scanned on a 3-T Siemens connectome-Skyra.

For each subject, four 15 minute fMRI scans with a temporal resolution of 0.73 seconds and a spatial resolution of 2-mm isotropic were available. The preprocessing pipeline followed the procedure outlined in Smith et al. [28, 29]. Spatial preprocessing was applied using the procedure described by [30]. ICA, followed by FMRIBs ICA-based X-noisefier (FIX) from the FMRIB Software Library (FSL) [31], was used for structured artifact removal, removing more than 99 percent of the artifactual ICA components in the dataset. Group spatial ICA was then used to obtain a parcellation of 50 components that cover both the cortical surfaces and the subcortical areas. Global signal regression was not employed. The parcellation was used to project the fMRI data into 50 time series.

The fMRI time series (size: number of participants × number of scans × number of time points × number of ICA components = 820 × 4 × 1200 × 50) were standardized so that, for each scan, subject, and ICA component, the data have a mean of 0 and SD of 1.

### 2.3 Simulation Study

To assess and compare the performance of the HMM versus the HSMM, we analyzed three different simulated data sets. The first data set was designed to evaluate the performance of each model when the true sojourn times are longer and the signal to noise ratio in the BOLD response variables is relatively large. The second was designed to evaluate the performance under the same sojourn times, but with a smaller signal to noise ratio. The third was designed to compare the two model choices when state switching times are very quick and the signal to noise ratio is large.

Each of the sets of simulated data consisted of 100 subjects, 1000 time points, 10 components, and 4 states. For each data set, the true sequence of states was drawn via both a transition probability matrix and each state’s sojourn distribution. The sojourn distributions in data sets one and two were Poisson distributed with λ = 20, 30, 40, and 50 for states 1,2,3 and 4 respectively. In the second data set, the sojourn distributions were geometric with p = 0.3, 0.4, 0.1 and 0.7. After the true state sequences were randomly drawn under these distributions, resting-state fMRI time series data was simulated from the four states using SimTB, a Matlab fMRI simulation toolbox [32]. The 4 states and 10 components were identical to those used in [6]. Gaussian noise was added to the simulated time series in data sets 1 and 3 with a standard deviation of 0.1, as was done in [6]. Additional noise was added to data set two (standard deviation of 0.6). See Figures 2-4 and Supplementary Figures S1 and S2 for the true sojourn distributions, transition probability matrices, and modular structure of each state.

To evaluate the accurateness of the HMM and the HSMM, we fit a HMM, a HSMM with a smoothed nonparametric sojourn density, and a HSMM with a Poisson sojourn density to each of the three simulated data sets. States, transition matrices, sojourn distributions, and empirical sojourn distributions were estimated under each model for each data set and compared to the truth. The state correlation matrices and the transition probability matrices were compared to the true states and transition probabilities by summing up the Euclidean distance between true and estimated matrix entries (i.e. correlation and transition matrices). We compared empirical sojourn distributions estimated after fitting the model to the data to the ‘true’ empirical sojourn distributions derived from the simulated data itself via the Kolmogorov-Smirnov test statistic. The Kolmogorov-Smirnov test is a non-parametric test of the equality of two probability distributions, and the Kolmogorov-Smirnov test statistic quantifies a distance between the empirical distribution functions. Section 3.1 presents these results.

### 2.4 Hidden Markov Modeling

We assume we have observed fMRI time series data denoted by *Y*_1_, …,*Y_T_*, where each vector *Y_t_* ∈ ℝ^1×*p*^ contains the BOLD measurements of *p* nodes at the *t^th^* time point. Throughout this paper, we assume that each *Y_t_* follows a multivariate Gaussian distribution 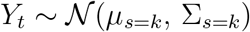, where the mean and covariance are dependent on the current network state *k* at time point *t*.

The collection of vectors of observed fMRI time series data *Y*_1_, …,*Y_T_* is denoted by *Ỹ*. We represent a latent/hidden network index variable underlying the observed time series vectors at a particular observation time by *S_t_*. The vector of latent segment variables, *S_1_,…, S_T_* is denoted by *S̃*. See Figure 1 for a visual representation. As is typical in a HMM, we make the following assumptions.

1. The vector of latent network states follows a first order Markov chain, i.e.

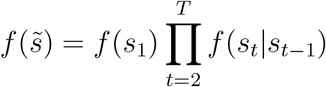
2. The observed BOLD signal vectors at each time point are conditionally independent given the latent process, i.e.

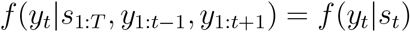

**Figure 1:**
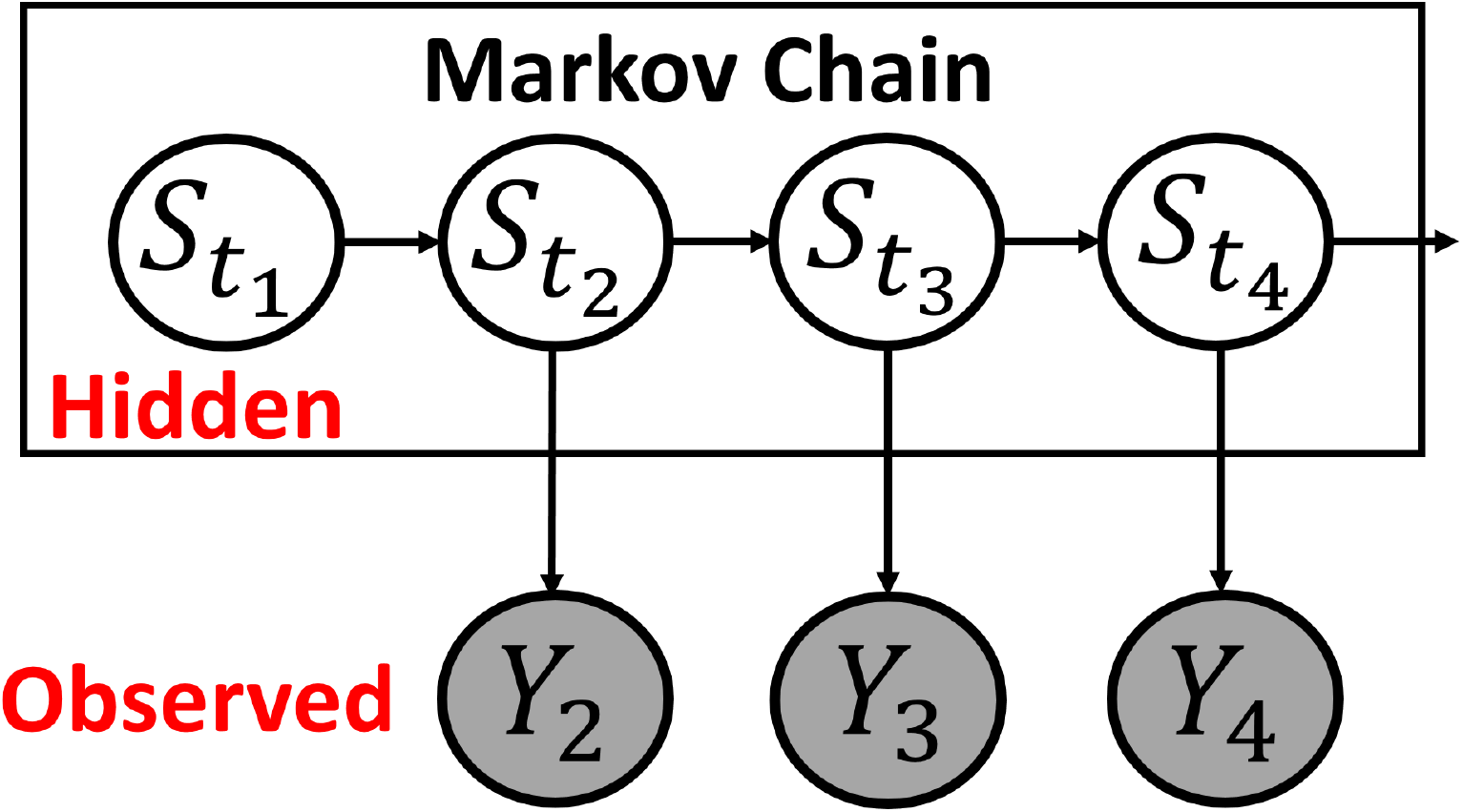
Hidden Markov Model Overview. The unobserved hidden states evolve according to a Markov process. The BOLD signals at each time point are assumed to follow a Gaussian distribution with parameters that depend on the current network state.

**Figure 2:**
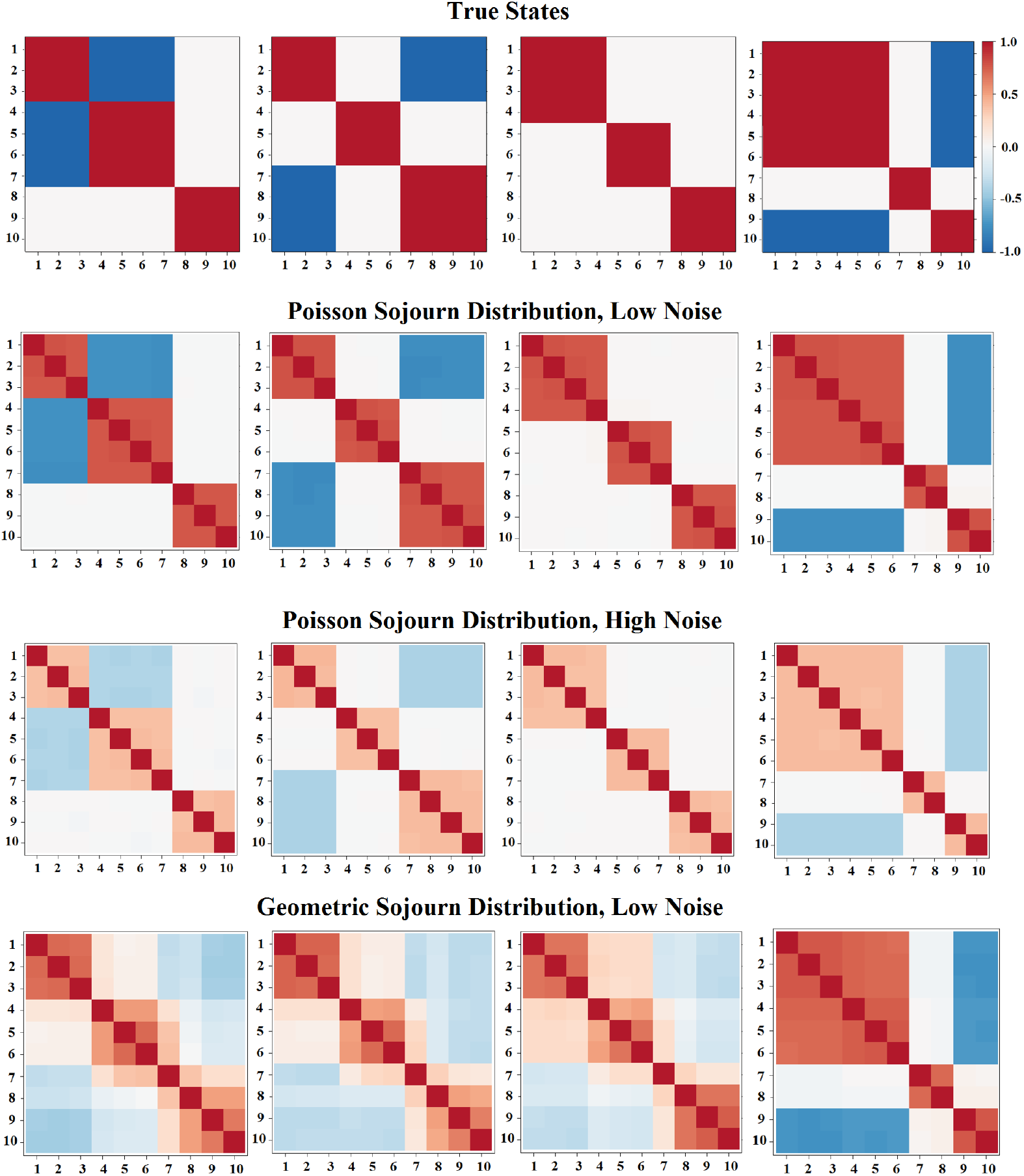
(Row 1) Four states from which fMRI data were simulated. (Row 2) States that were estimated via the HMM and HSMMs under the simulation scenario of Poisson sojourn distributions and low noise. (Row 3) States that were estimated via the HMM and HSMMs under the simulation scenario of Poisson sojourn distributions and high noise. (Row 4) States that were estimated via the HMM and HSMMs under the simulation scenario of Geometric sojourn distributions and low noise. Only one set of estimates states is shown for each simulated data scenario since all three models produced nearly identical results.

Under these assumptions, the complete data log-likelihood for a single subject, assuming there are *K* network states, can be written

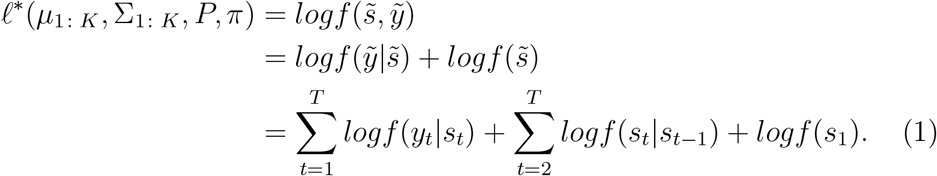

The first term is based on the conditional distribution of the observed BOLD signal vector given the underlying *k^th^* hidden network, which takes on a Gaussian distribution:

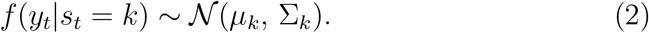

The second term is based on a transition probability matrix, denoted *P*, where the probability at the *i^th^* row and *j^th^* column represents the probability of transitioning from network state *i* to state *j* (i.e. *p_ij_* = *P*(*S_t_ = j*|*S*_*t*−1_ = *i*)). The third term is the distribution of the network state at the first time point. In practice this is typically represented as a vector *π*.

### 2.5 Hidden Semi-Markov Modeling

An extension to the HMM is the hidden semi-Markov model (HSMM), which allows the underlying latent process to be a semi-Markov chain. In a standard HMM, the sojourn time (i.e. number of consecutive time points in a state) distribution for a given state is implicitly geometrically distributed [24]. To see why this is true, consider the probability of spending u consecutive time points in state *k* once state *k* is entered, which is given by

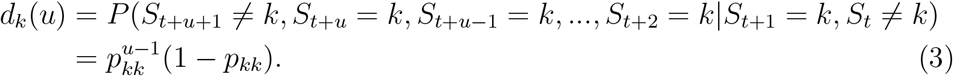

This implies that the probability of remaining in a given state decreases as the sojourn time increases. Therefore, more weight is placed on shorter consecutive time points in a given state. We hypothesize that this may not always be appropriate in the context of FC analysis.

The HSMM differs from the standard model in that the sojourn time is explicitly defined in the model, and therefore, able to be estimated. Ferguson [33] was the first to propose such models. If one has prior knowledge as to what family of distributions the sojourn density may follow (e.g. a Gamma or Poisson distribution), one could parameterize the sojourn specification of the likelihood accordingly. Otherwise, one may choose to fit a nonparametric distribution to the sojourn time and let the data fully drive the density estimate. An incorrectly specified sojourn density has the ability to impact the results in a negative way by leading investigators to believe subjects are switching states far too often. Moreover, the sojourn density may be an important piece of information for better understanding disease mechanisms. If this is the case, it is important to have an accurate estimate, instead of one forced to be geometrically distributed.

The complete data log-likelihood of the HSMM for a single subject, assuming there are *K* unique network states, can be written as

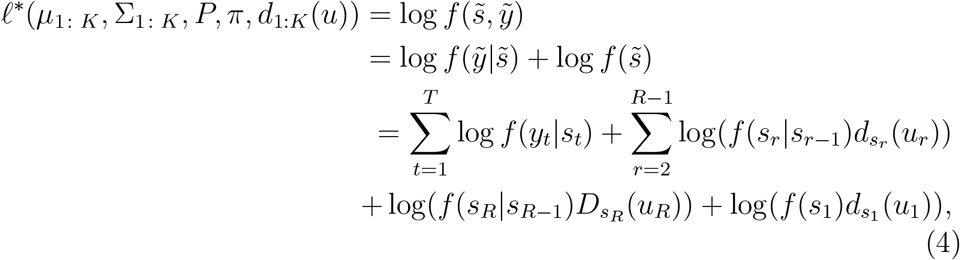

where *s_r_* is the *r^th^* visited state, *u_r_* is the number of consecutive time points spent in that state, and *d*_*s*_1__ (*u*_1_) is the sojourn density for the first entered state. Note this model has an extra term for the sojourn distribution, *d_s_r__*(*u_r_*), which was not present in (1) for the standard HMM. Also note that

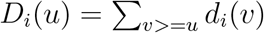

is the survivor function and pertains only to the sojourn time in the final state. Developed by Guedon [34], it allows us to *not* make the assumption that the process is leaving the final state immediately after time *T*.

### 2.6 Maximum Likelihood Estimation for the HMM and HSMM

The parameters of both models were estimated utilizing the *mhsmm* package for R [35]. This package allows for inference for multiple observation sequences (i.e., multiple subjects at once). The Expectation-Maximization algorithm [36], an iterative method which alternates between performing an expectation (E) step and a maximization (M) step, was used to estimate all model parameters. First, in the E-step, the expected value of the complete data log-likelihood with respect to the unknown network states *s̃*, given the observed BOLD series vectors *ỹ* and the current parameter estimates, is calculated. This calculation involves estimating several terms: (1) The probability of being in state *k* at time *t* given the observed sequence of BOLD signals,

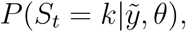

where *θ* is the collection of current parameter values, (2) the probability that a subject leaves state *i* at time *t* and enters state *j* at time *t* + 1 given the observed sequence,

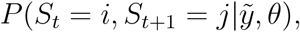

and in the HSMM, we also need the expected number of times a subject spends *u* time steps in state *k*. These quantities are calculated via a well-known dynamic programming method known as the forward-backward algorithm. Please see [37] and [34] for details regarding this algorithm.

The expected log-likelihood found in the E-step is then maximized in the M-step. The initial state probabilities, transition probabilities, and means and covariances for each state are estimated via closed form solutions. See [35] for the formulas for these. In the HSMM, the state duration density must also be estimated. For example, for a shifted Poisson distribution, λ_*k*_ is estimated for all possible shift parameters d, choosing the *d* which gives the maximum likelihood. Again, please refer to [35] for additional details. The E- and M-steps are repeated until convergence.

The computation time of the above algorithm is a function of the number of subjects, number of time points, number of ROIs, number of states, and number of model parameters in the sojourn density. As an example, the anxiety data set described in Section 2.1 (23 subjects, 215 time points, and 5 ROIs) took less than a minute of computing time to estimate the model parameters. However, the HCP data set described in Section 2.2 (1200 subjects, 4 scans, 1200 time points for each scan, and 50 ROIs) took 1-2 weeks to estimate the model parameters.

### 2.7 HSMM Sojourn Distribution Specification

In this paper, we used two different sojourn distributions when working with HSMMs; a shifted Poisson distribution and a smoothed nonparametric distribution. The shifted Poisson distribution takes the form:

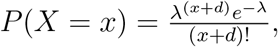

where *d* is the shift parameter. The smoothed-nonparametric sojourn distribution is particularly useful for situations when we don’t know the exact form of the sojourn distribution, which is especially true when analyzing resting-state fMRI. It does not have an analytic form. We initialized the model with a uniform distribution where the parameters are a *T × K* matrix (*T* represents the total number of time points and *K* the number of states). For each state, at each iteration of the E-M algorithm, a kernel density estimate was computed by first finding which *t* in the range [1, *T*] has a density value that exceeds a smoothing threshold (1*e*^−20^) and then by using the density value at those points together with a Gaussian kernel (for smoothing purposes) to obtain the estimate.

### 2.8 Choosing an Optimal Number of States

A key challenge when working with either HMMs or HSMMs is determining the optimal number of states to include in the analysis. For the anxiety-inducing data set in this work, we employed a cross-validated BIC criteria to choose the number of states. For each choice of *k* = 1, … *K*, we performed the following steps:

1. For *i* = 1, …, *n*, where *n* is the number of subjects:

a. Fit the HSMM (or HMM) on the remaining *n* − 1 subjects via the E-M algorithm, and save the parameter estimates.
b. Calculate the leave-one-out log-likelihood for the *i^th^* observation as follows:

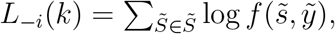

where the summation is over all possible hidden network sequences, *f*(*s̃*, *ỹ*) is defined as in (1) or (4), and the estimated parameters from (a) are used.
2. Calculate

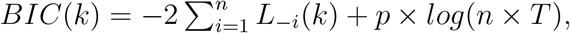

where *p* is the number of model parameters and *T* the number of time points in the scan.

After performing these steps, we chose the value of *k* that minimized the BIC criterion in step 2.

### 2.9 State Matching

When comparing the results between the standard HMM and the HSMM with different sojourn distribution specifications (i.e., Poisson or nonpara-metric), it is necessary to match the state labels. To do so, we employed an algorithm based on computing the Euclidean distance between the covariance matrices associated with each state.

We first chose a set of states to be considered our reference states. For example, in the HCP study, the states inferred from fitting the HSMM with the nonparametric sojourn distribution on all of the data were our reference states. We labeled these states {1, 2, …, *K*}. We then took the *K* states inferred from the model we want to match to the reference and labeled them {1*, 2*, …, *K*\*}. The Euclidean distance was then calculated between all pairs of states between the two groups.

We first found the best match in the set {1*, 2*, …, *K*\*} for each state in {1, 2, …, *K*}. Next, we reversed the order and find the best match in the set {1, 2, …, *K*} for each state in {1*, 2*, …, *K*\*}. We deemed final matches for any pairs where the two directions were in agreement. Any pairs that were not in agreement were grouped into a single pool and re-matched with each other so that the total sum of the Euclidean distance between all pairs was minimized.

### 2.10 Multi-dimensional Scaling

Multidimensional scaling takes a set of dissimilarities as input and returns a set of coordinates such that the distances between the coordinates are approximately equal to the dissimilarities. We used this technique to visualize the distance between different states on a lower-dimensional space. Euclidean distance is used as the dissimilarity measure and was calculated between all pairs of states inferred from the HSMM. The *CMDSCALE* function in R, which follows the analysis procedure outlined in [38], was used to calculate the best-fitting two-dimensional representation of the states.

### 2.11 Defining Significant Edges

It is often useful to infer a binary graph associated with each state (such as those presented in Figure 5), as opposed to simply looking at the covariance/correlation matrices. There are various ways to infer such networks, but it is sometimes desirable to construct a graph *G* where the inferred edges are representative of a direct relationship among vertices, rather than an indirect relationship. In this case, partial correlation is often preferred.

In the context of FC where we assume the BOLD signals at each ROI follow a multivariate Gaussian distribution, partial correlation measures the degree of association between each pair of ROIs, while controlling for the effects of all other ROIs. Under the Gaussian assumption, and denoting the partial correlation coefficients as *ρ_ij_|V*\{*i,j*}, vertices *i, j* ∈ *V* have *ρ_ij_|V*\{*i,j*} = 0 if and only if the BOLD signals at regions *i* and *j* are conditionally independent given the BOLD signals at all other regions. The overall model, combining the multivariate Gaussian distribution with the graph generated by defining edges for all pairs of vertices where *ρ_ij_|V*\{*i,j*} = 0, is called a Gaussian graphical model [39].

In the context of Gaussian graphical models, partial correlation coefficients can be expressed in the form:

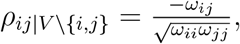

where *ω_ij_* is the (*i,j*)^*th*^ entry of Ω = Σ^−1^, the inverse of the covariance matrix Σ. The covariance matrices are estimated for each of the *K* states in the HMM or HSMM.

We utilized a method suggested by Drton and Perlman [40] that addresses the problem of testing which pairs of partial correlations are not equal to 0, as well as the multiple testing problem of having to perform a separate test for each pair of vertices. After first Fisher-transforming the partial correlations using the formula

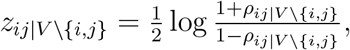

we assigned an edge to each vertex pair *i, j*, if and only if,

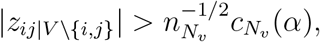

where *n* is the number of subjects, *N_υ_* is the number of vertices, *n_N_υ__* = *n−N_υ_*, and

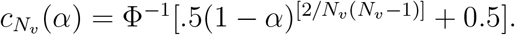

Here Φ^−1^ is the cumulative distribution function of the Gaussian distribution with mean 0 and variance 1. If we assume that *G* is the true conditional independence graph underlying our data, then it can be shown that this procedure correctly infers *G* with probability at least 1 − *α*. It is important to note that this procedure is not applicable when the number of vertices exceeds the number of subjects.

## 3 Results

### 3.1 Simulation Study

A standard HMM, a HSMM with a nonparametric sojourn density, and a HSMM with a Poisson sojourn density were fit to the three simulated data sets described in Section 2.3, with the goal of assessing how accurate each model is in estimating the true transition matrix, sojourn densities, and states. Figure 2 presents the four true states from which resting-state fMRI data were simulated in all three data sets, as well as the states estimated from each data set. In all three data sets, the results were essentially identical across all three models fit, leading us to present one set of estimated states for each data set. Both the HSMMs and standard HMM do an excellent job of estimating the states from the data sets simulated with a Poisson sojourn density. The Euclidean distances between the true and estimated states are 1.65, 1.55, 1.13, and 1.76 for states 1-4 respectively from the HMM model; 1.66, 1.56, 1.13, and 1.76 from the HSMM with a nonparametric sojourn density; and 1.66, 1.56, 1.13, and 1.77 from the HSMM with a Poisson sojourn density. The data set with more noise has smaller absolute values of the correlations, which is to be expected with noisier data. However, the modular structure of the states is still well defined. The Euclidean distances between the true and estimated states are 4.27, 4.14, 2.97, and 4.59 for states 1-4 respectively from the HMM model; 4.27, 4.16, 2.98, and 4.61 from the HSMM with a nonparametric sojourn density; and 4.27, 4.17, 2.99, and 4.62 from the HSMM with a Poisson sojourn density. Interestingly, however, none of the three models do a great job with estimating states from simulated data that arose from sojourn times being geometrically distributed. The estimate of state 4 is the most accurate, while the other three state estimates appear to be a mix of one another. The Euclidean distances between the true and estimated states are 5.74, 4.58, 3.35, and 2.21 for states 1-4 respectively from the HMM model; 5.57, 4.43, 3.33, and 2.23 from the HSMM with a nonparametric sojourn density; and 5.85, 4.53, 3.37, and 2.11 from the HSMM with a Poisson sojourn density. Notably, state 4 had the slowest rate of decay in its sojourn density. In other words, simulated subjects spent more consecutive time points in state 4, compared to the other three states. This suggests the possibility that neither model performs well when estimating states from data where subjects are switching very rapidly from state to state.

Figure 3 presents the true transition matrices for each data set, as well as the estimated transition matrices for each model fit on each data set. The top row of the figure suggests that all three models perform fairly well in the case of a Poisson sojourn density and relatively low noise. However, the transition matrix estimated from the HMM is somewhat less accurate compared to the HSMMs. The Euclidean distances between the true and estimated transition probability matrices are 0.20, 0.11, and 0.076 for the standard HMM, HSMM with nonparametric sojourn density, and HSMM with Poisson sojourn density, respectively. The pattern of positive and negative correlations is accurate, but the strength of the correlations are not quite as they should be. We see the same behavior in the higher noise case, as well, with the Euclidean distances being 0.24, 0.10, and 0.072. Moreover, just as in the estimation of states, the data simulated via a HMM does not lead to accurate estimations of the transition matrix with any of the models fit, with distances of 0.77, 0.63, and 0.63.

**Figure 3:**
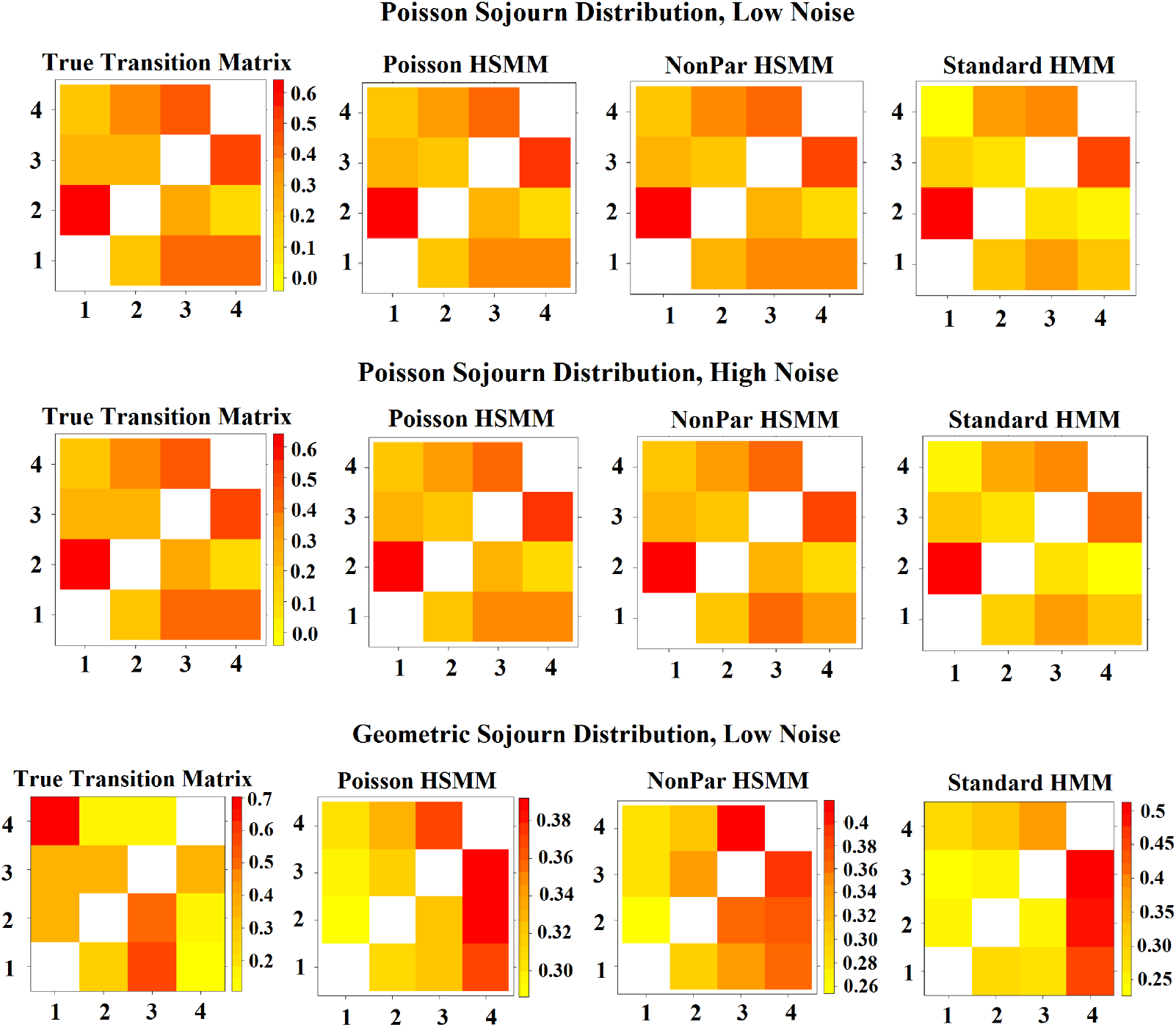
(Row 1) Transition matrices that were estimated via the HMM and HSMMs under the simulation scenario of Poisson sojourn distributions and low noise. (Row 2) Transition matrices that were estimated via the HMM and HSMMs under the simulation scenario of Poisson sojourn distributions and high noise. (Row 3) Transition matrices that were estimated via the HMM and HSMMs under the simulation scenario of Geometric sojourn distributions and low noise.

Figure 4 presents the sojourn distributions estimated from the Poisson distributed sojourn time data set with low noise data after fitting each of the three models, as well as the empirical sojourn distributions obtained after fitting the models and the empirical distributions estimated directly from the simulated data. Figures S1 and S2 in the supplement present the same but for the remaining two simulated data sets. Quite expectedly, the HSMM with a Poisson sojourn density does well with estimating the sojourn distributions from the data sets simulated from a Poisson sojourn density (both low and high noise). The HSMM with a smoothed-nonparametric sojourn density also estimates the true Poisson sojourn densities well. However, as we suspected, fitting the HMM leads to sojourn distributions that place too much weight on quicker state switches. The sum of the Kolmogorov-Smirnov test statistics across the four states for the comparison of the ‘true’ empirical sojourn distributions calculated directly from the Poisson simulated data set with low noise to the empirical estimate obtained after fitting each of three models to the data set are 0.84, 0.76, and 0.36 for the standard HMM, HSMM with nonparametric sojourn density, and HSMM with Poisson sojourn density, respectively. For the lower noise data set, the test statistics are 1.20, 1.01, and 0.56, respectively. In both cases, the HMM performs the worst. Lastly, when fitting the three models on the data derived from geometric distributions, the HSMM with the nonparametric sojourn density and the standard HMM appear to perform similarly with regards to the empirical sojourn distributions, suggesting that the HSMM may be capable of estimating quick state switches just as well as the standard HMM. The sum of the Kolmogorov-Smirnov test statistics across the four states are 1.322, 1.309, and 1.458 for the standard HMM, HSMM with nonparametric sojourn density, and HSMM with Poisson sojourn density, respectively.

**Figure 4:**
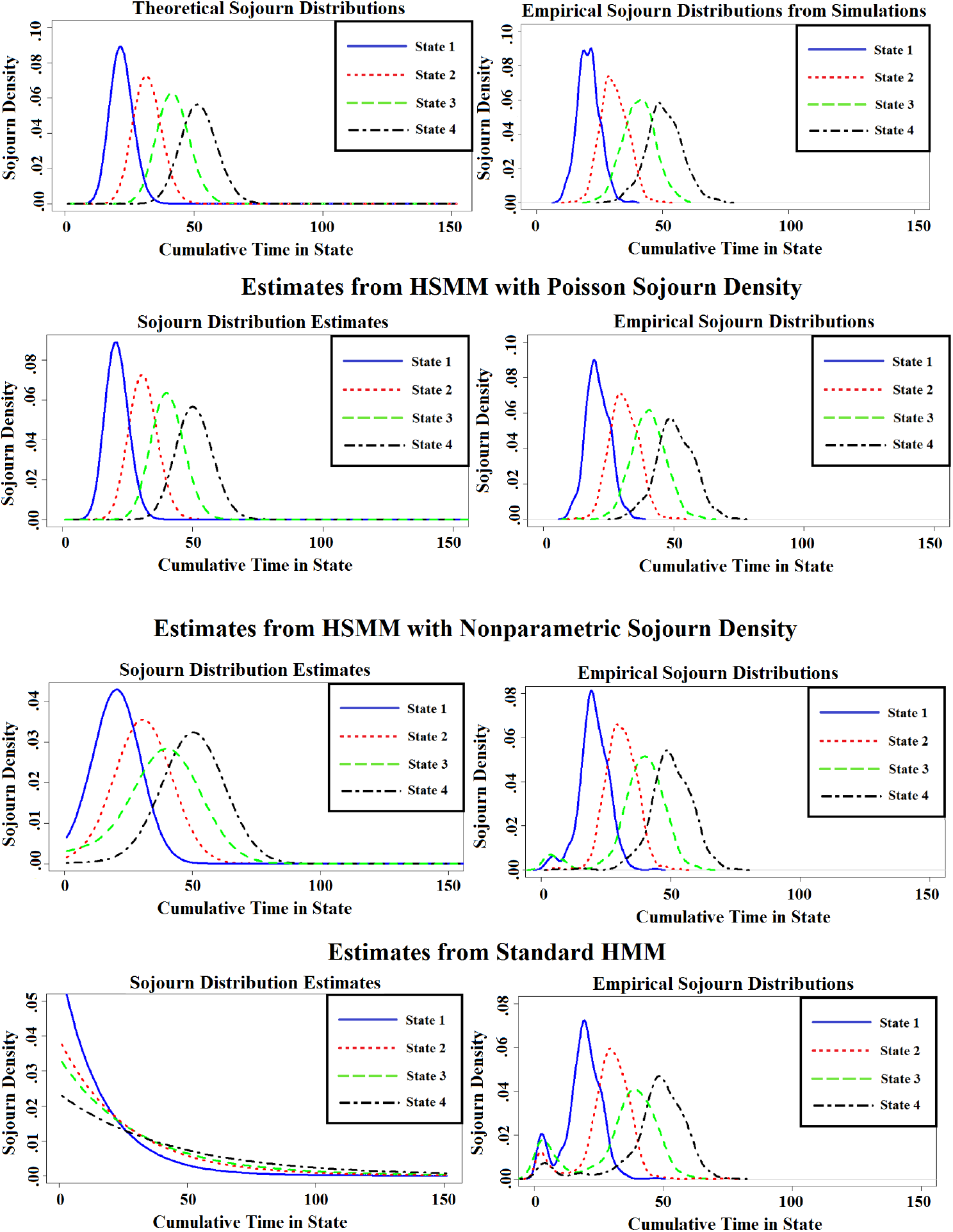
Sojourn distributions, as well as empirical sojourn distributions, that were estimated via the HMM and HSMMs under the simulation scenario of Poisson sojourn distributions and low noise.

While an extensive simulation study is required for a more in-depth comparison between these two flavors of models in many different settings, these initial results suggest that if the goal is to estimate the states and transition matrices, either model may be fine. However, if the goal is to estimate state sequences and sojourn times, the HSMM outperforms the HMM when sojourn times do not truly follow a geometric distribution. In the case where true sojourn times do follow a geometric distribution, the HSMM with a nonparametic sojourn density appears to perform just as well as the standard HMM when estimating sojourn times. These results demonstrate the flexibility of the HSMM and the potential inaccuracy of the standard HMM when true sojourn times are unknown or are believed to not be geometrically distributed. As more researchers are moving towards analyzing transition times for clinical populations, we feel it is important to obtain as accurate an estimate as possible of the state sequences and sojourn times.

Lastly, if a state’s sojourn distribution is truly geometric (which we feel is not common with fMRI data), one may need to proceed with caution when estimating the states and transition matrices. We leave further evaluation of the performance of these models to future work.

### 3.2 Anxiety Inducing Experiment

We analyzed the fMRI data from the 23 participants who performed the anxiety-inducing task (*Methods Section 2.1*). The data consisted of 4 brain regions and a dummy region corresponding to heart rate. Figure 5 depicts the experimental paradigm.

**Figure 5:**
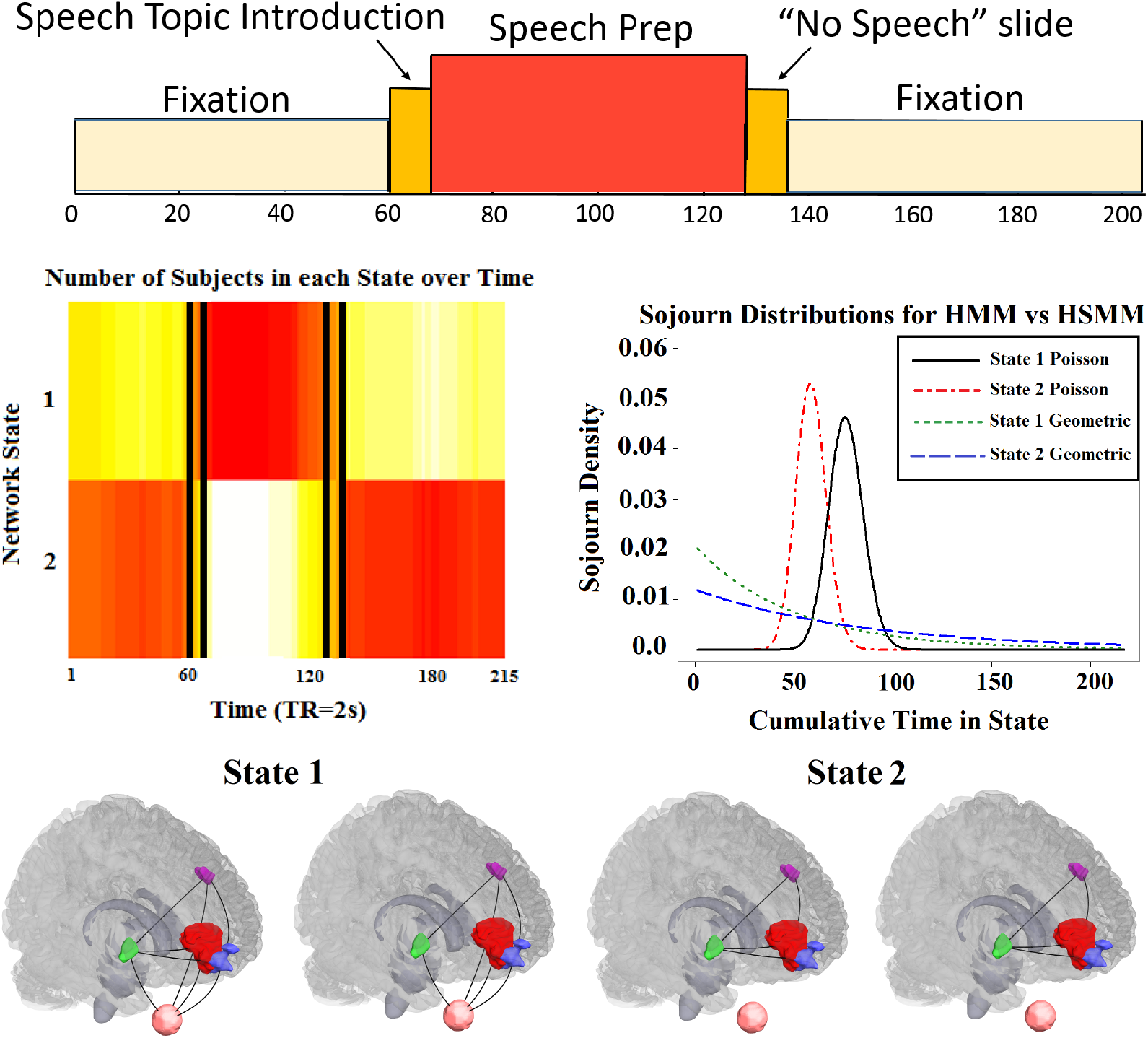
(Top row) Experimental paradigm for the anxiety-inducing task [27]. (Row 2 left) Heatmap demonstrating that the HSMM closely uncovers the design of the experiment. Vertical bars indicate the true design. (Row 2 right) Sojourn distributions under both a standard HMM and a HSMM for both network states. The Poisson distributions estimated under the HSMM closely match the experimental design, while the geometric distributions, implicit in the HMM and parameterized by the diagonal of the transition matrix, do not. (Row 3) States 1 and 2 estimated via the HSMM for two different significance levels (α = 0.05 and α = 0.01, respectively for each state). Regions consist of two different subsections of the ventral medial prefrontal cortex (red and blue), the dorsal lateral prefrontal cortex (purple), and the ventral striatum (green), Heart rate (pink) is depicted below each brain image.

#### 3.2.1. HSMM Uncovers Task Alignment Near Perfectly with Two-State HSMM

A HSMM was fit assuming two network states and a shifted Poisson sojourn distribution (*Methods Sections 2.5-7*). The network states inferred by the HSMM represent unique FC graphs of the 5 regions that repeat at different points in time. Each subject’s most probable series of true network states was estimated using the Viterbi Algorithm, a well-known dynamic programming algorithm used to find the most likely series of hidden states, given a series of observed states [41].

A heatmap presenting the number of subjects in each state at each time point of the scan is shown in Figure 5. The heatmap depicts a clear pattern where the majority of participants begin in State 2 during the fixation period, transition to State 1 during the speech instruction slide and speech preparation period and then return to State 2 during the final fixation period. Clearly, even though the HSMM is agnostic to the experimental paradigm, it still discovered the alignment near perfectly.

The estimated network states are shown in the bottom row of Figure 5 in the form of binary graphs. Networks were estimated by inverting the estimated covariance matrices, calculating partial correlation matrices, and thresholding correlations to create binary edges based on a method for estimating Gaussian concentration graphs (for two different significance levels)(*Methods Section 2.11*). Of particular interest is that during the high anxiety state (State 1), heart rate is associated with all four brain regions, while in the low anxiety (baseline) state, the heart rate related edges equal 0.

The sojourn distributions for both network states under the HSMM are also shown in Figure 5. For comparison, we fit a standard HMM to the data and plotted the sojourn distributions implicit in this model. The Poisson distributions estimated under the HSMM closely match the experimental design. For example, the mean sojourn time for state 2 is approximately 60 time points (i.e. 2 mins), which is a reasonable estimate given the pattern that emerged in the heatmap. The geometric distribution, however, does not appear to fit the data well, as it indicates that subjects have a high probability of spending only a few seconds in either one of the states before immediately switching to the other state. Thus, the incorrectly specified sojourn distribution places more weight on short amounts of consecutive time in a state.

#### 3.2.2. HSMM and HMM produce different results

Next, two HSMMs were fit assuming three network states, as three was the number that minimized our cross-validated model selection criterion in the nonparametric sojourn distribution setting (*Methods Section 2.8*). We fit the HSMM with a shifted Poisson sojourn distribution, as well as a smoothed-nonparametric sojourn distribution (*Methods Section 2.7*). For comparison, we also fit a standard HMM. Results are shown in Figure S3 in the supplement.

The row of heatmaps at the top of the figure reveal the number of subjects estimated to be in each of the three states at each time point. Differences between the three models are somewhat subtle, but they do exist. The HSMM fit with a Poisson sojourn density appears a bit ‘smoother’ in the sense that the colors are more blended throughout. The smoothness of the Poisson sojourn HSMM results is also evidenced by the second row of plots. These plots show the estimated sequence of states for each subject, where each color represents a different state. As can be seen by the plots, there are fewer state switches in the HSMM with a Poisson sojourn density.

Lastly, the bottom row of Figure S3 shows the estimated sojourn distributions for each state under the three different models. The standard HMM places more weight on quicker states switches, whereas the Poisson sojourn distribution HSMM places the least amount of weight on quicker switches. This was reflected in the heatmaps and plots of the individual state sequences. However, given the large signal of the high anxiety provoking part of the experiment compared to the lower anxiety period, we shouldn’t expect to see a huge effect of a misspecified sojourn distribution. In other studies where the sojourn distribution is misspecified and the signal to noise ratio is smaller we anticipate it may be more costly to fit the standard HMM.

The estimated Pearson correlation matrices, representing the network states, from each model are presented in Figure S4 in the supplement. Previously, in the two-state model, we presented binary networks based on partial correlation. Here we present just the Pearson correlation matrices to demonstrate that there are a variety of ways one could choose to visualize the network states. In this depiction, the states estimated under the standard HMM and smoothed-nonparametric sojourn HSMM are nearly identical. The HSMM fit with a Poisson sojourn density has some differences. Most notably, in state 3, heart rate appears more negatively correlated with the ventral medial prefrontal cortex and dorsal lateral prefrontal cortex. However, these specific correlations are not significantly different from 0 in any of the three models.

### 3.3 Human Connectome Project Data

We next performed a dynamic FC analysis using the resting-state data from the HCP [28]. Data consisted of 50 brain regions of interest that cover both the cortical and subcortical areas obtained via group spatial ICA (*Methods Section 2.2*). Subjects were each scanned 4 times, where each scan contained BOLD signals at 1200 time points. We concatenated all subjects and scans, and we fit three separate models to the data, i.e. the standard HMM, the HSMM with a smoothed-nonparametric sojourn distribution, and the HSMM with a shifted Poisson sojourn distribution. Rather than use the BIC-criteria to determine the optimal number of states, we instead inferred 12 network states with all three models, as this was the number of states Vidaurre et al. [21] used in a recent standard HMM analysis using the same data.

#### 3.3.1. Transition Matrices Agree with Previous Findings

Despite different methods of estimation (Vidaurre et al. [21] utilized Variational Bayes, while we use the E-M algorithm), we first sought to replicate their findings. A main finding is the discovery that the dynamic switching between the 12 states is not random. They discovered two larger metastates while looking at the transition probability matrix, indicating that the brain has a tendency to cycle most often between network states within a metastate and less often between networks in different metastates.

We found similar results not only when using the traditional HMM, as they did, but also when we fit the two HSMMs. Figure 6 shows the estimated transition probability matrices obtained from fitting the models and matching the states from each model (*Methods Section 2.9*). In each of the three models, a pattern emerges where the states labeled in the upper left hand quadrant are more likely to be transitioned between, the states in the lower right hand corner are more likely to be transitioned between, and transitions between quadrants are less likely.

**Figure 6:**
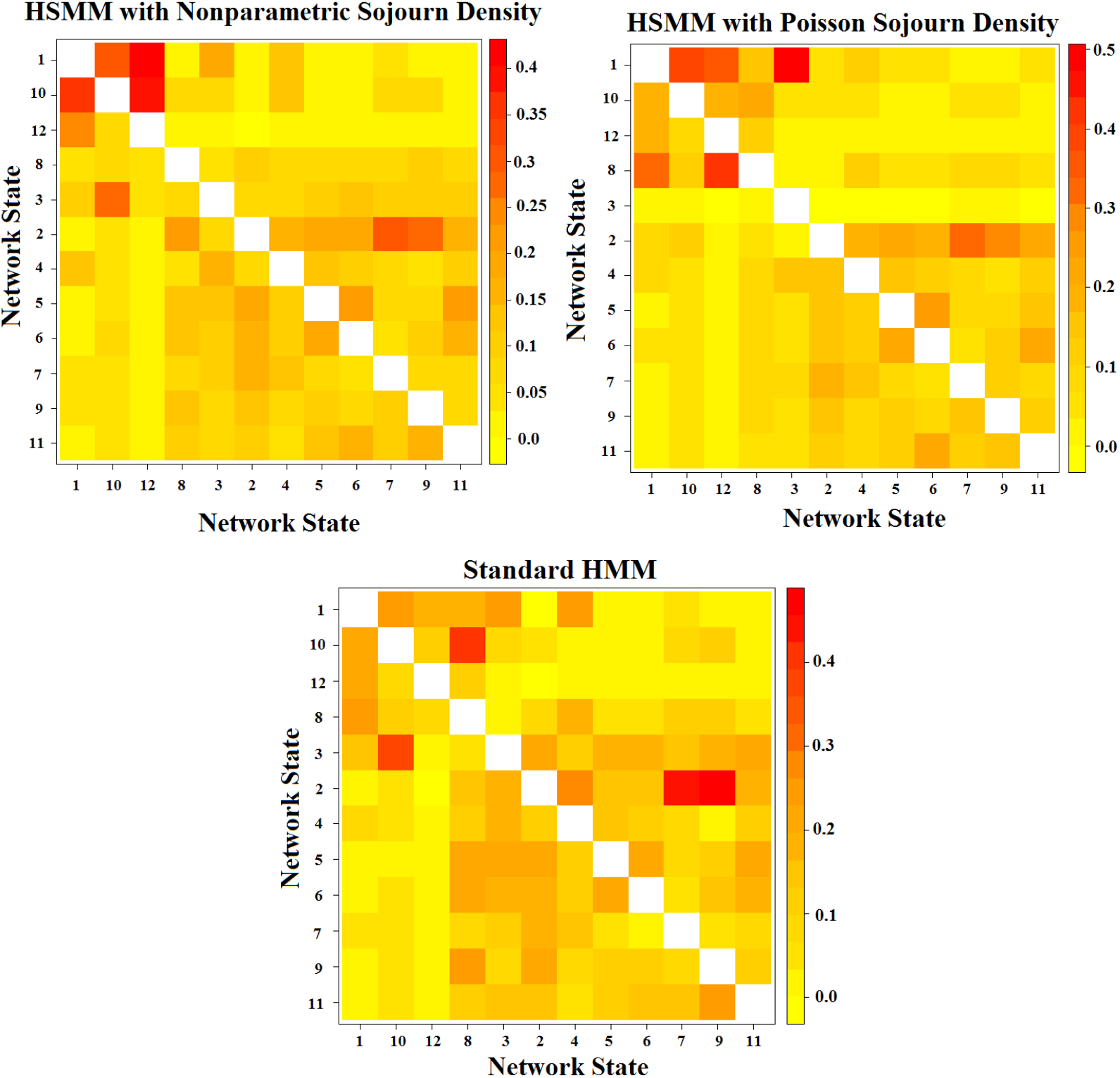
Transition matrices depicting the probabilities of transitioning between states from HCP data. (Top) Transition matrices for the two HSMM models. Probabilities are conditional on switching to a new state. (Bottom) Transition matrix for the standard HMM. Probabilities are not conditional on switching.

#### 3.3.2. Multidimensional Scaling Reveals Resting-State FC States are Reproducible

Next, we sought to determine how reproducible the network states are across both subjects and scans. To answer this question, we split the data set in various ways. First we separated the data by taking all scans for the first half of the subjects and then all scans for the second half of the subjects. We fit a separate HSMM with the smoothed-nonparametric sojourn distribution to each of these groups. Then, we split the data by scan. We took scans 1 and 2 for all subjects and separated them from scans 3 and 4 for the same subjects. Again, we fit a separate HSMM with the smoothed-nonparametric sojourn distribution to each of these groups.

The 12 states inferred for each of the above sub-analyses were collected. All states were matched to the original 12 states inferred using the complete data set. Multidimensional scaling was performed for all sets of states. The results are presented in Figure S5 in the supplementary material. In each plot, the distances between the coordinates are approximately equal to the dissimilarities of the states (i.e. covariance matrices) (*Methods Section 2.10*).

The pattern in all 5 plots are extremely similar. States 1, 4, 8, 10 and 12 are separated from the other states. Interesting enough, four of these five states belong to the top left quadrant of the estimated transition probability matrix, indicating that they are often cycled between. States 7 and 9 also appear to be relatively close to each other. Lastly, the remaining states are clustered together in one group. These results demonstrate that the 12 states are a reproducible and robust estimation of brain activity in the population.

Figure 7 is an alternative representation of the multidimensional scaling plot created for the 12 states inferred from the entire dataset. The states are clustered according to distance, with some states in their own cluster. The width of the arrows are proportional to the probabilities of transitioning to and from each cluster. As was also indicated by the transition probability matrix, states 1, 10 and 12 have high transition probabilities to and from each other. The asterisks assigned to states 2, 5, 6, and 11, indicating that they have higher probabilities of transitioning within their cluster than to states outside of their cluster, match our conclusions from the transition probability matrix. In general, the figure demonstrates that the clusters that are closer together typically have higher probabilities of transitioning between each other, indicating that the probability of transitioning between states is influenced by the similarity in functional connections.

**Figure 7:**
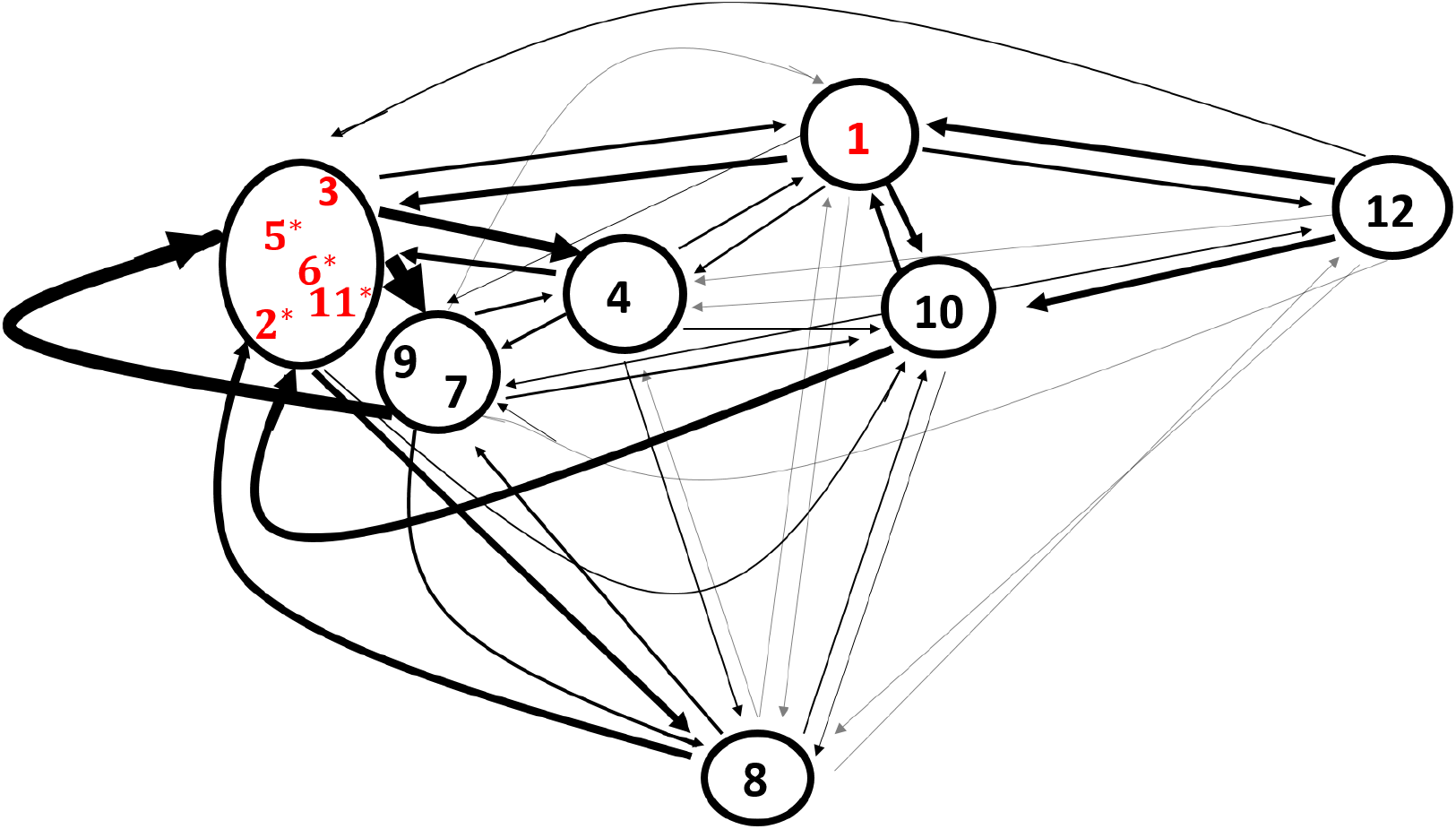
Multidimensional scaling representation of the 12 network states. Relative position of one state compared to another state is determined by the Euclidean distance between covariance matrices. Edge widths are proportional to the transitional probabilities between states, with thicker edges representing a higher probability. Edges corresponding to a transition probability < 0.05 are lightened and any with a probability < 0.01 are not included. Red state labels indicate that the state was one of the top 6 states in terms of proportion of time spent in the state. Asterisks highlight states where the probability of transitioning between states within the same cluster is greater than the probability of transitioning to states outside of the cluster.

States 2 and 12 are shown in Figure 8 and represent two states that are quite different from each other (as indicated by their distance in the multidimensional scaling plot). We present both the Pearson and partial correlation matrices, as both are popular metrics for estimating connectivity between nodes. However, we interpret only the Pearson correlation matrices in this case. State 2 appears to have a majority of connections within dorsal and ventral attention regions and also within DMN regions. State 12 appears to have fewer connections within the DMN and more connections both within the cerebellum and within the visual regions. There also appear to be strong connections between ventral attention regions and somatomotor regions. The remaining ten states are presented in the supplementary material.

**Figure 8:**
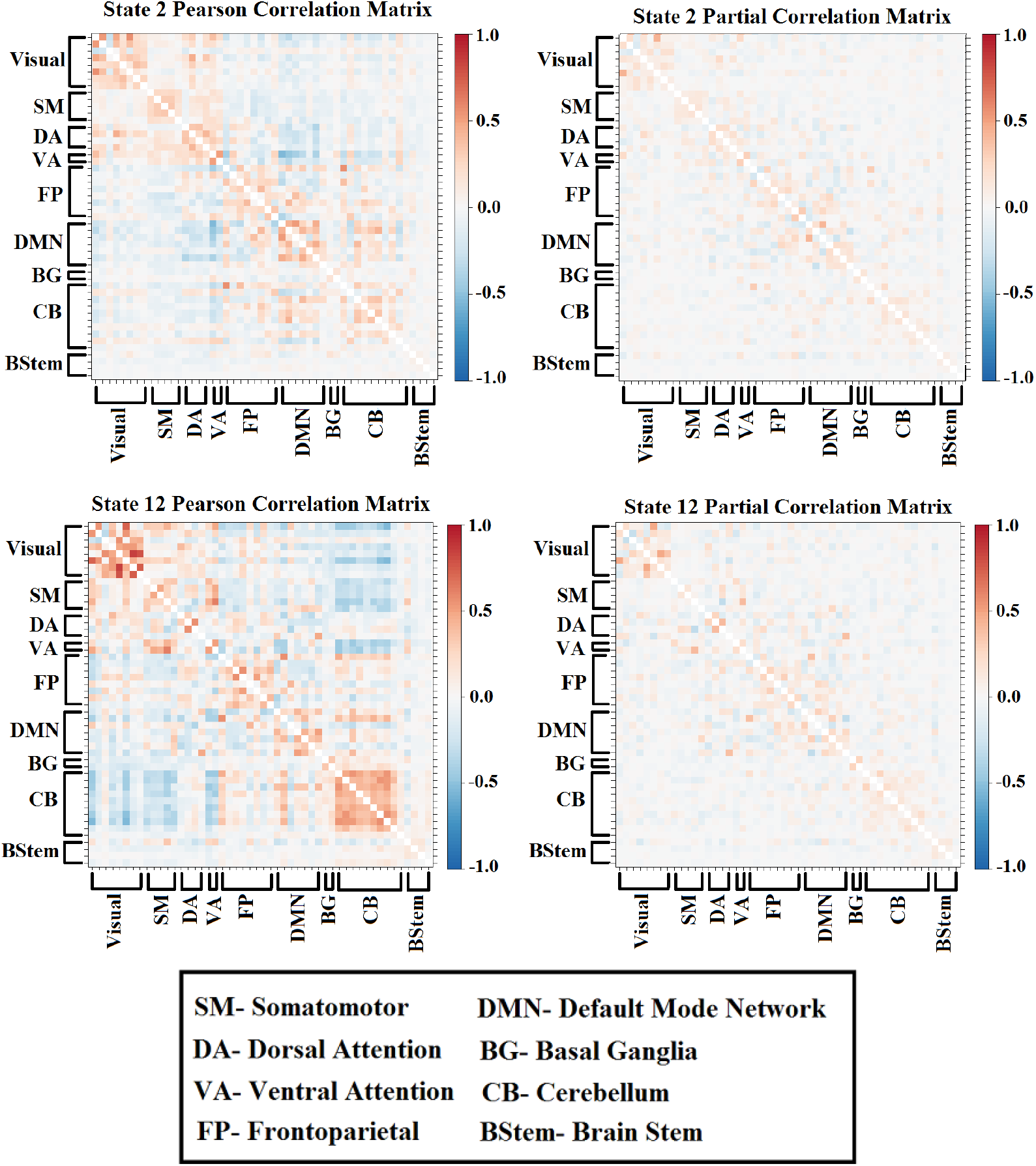
Pearson and partial correlation matrices for two of the states inferred by the HSMM with the smoothed-nonparametric sojourn distribution. State 2 has connections within dorsal and ventral attention regions and also within default mode network regions. State 12 has connections both within the cerebellum and within the visual regions. Moreover, there are strong connections between ventral attention regions and somatomotor regions.

#### 3.3.3. Assumed Geometric Sojourn Distributions in the Standard HMM have an effect on Empirical Sojourn Times

Although the estimated transition probability matrix is similar between the standard HMM and the two different HSMMs, subject sojourn times are influenced by the model choice. Figure 9 gives a comparison of the estimated sojourn distributions for the two states where subjects spent the greatest proportion of time (i.e. states 2 and 5). The sojourn distributions estimated directly from the standard HMM and the HSMM with a nonparametric sojourn distribution are shown in bold print. The dotted curves represent the empirical sojourn distributions under each model. Empirical sojourn distributions were calculated by estimating each subject’s most probable state sequence for each of his/her scans, tallying the sojourn times across subjects and scans, and then estimating a smooth density curve with a Gaussian kernel. As is shown by the plot, the standard HMM places more weight on quicker state switches. In practice, this has an influence on the sojourn times estimated across subjects, as indicated by a difference in empirical sojourn distributions. The empirical distribution estimated under the HMM is pulled towards the geometric distribution.

**Figure 9:**
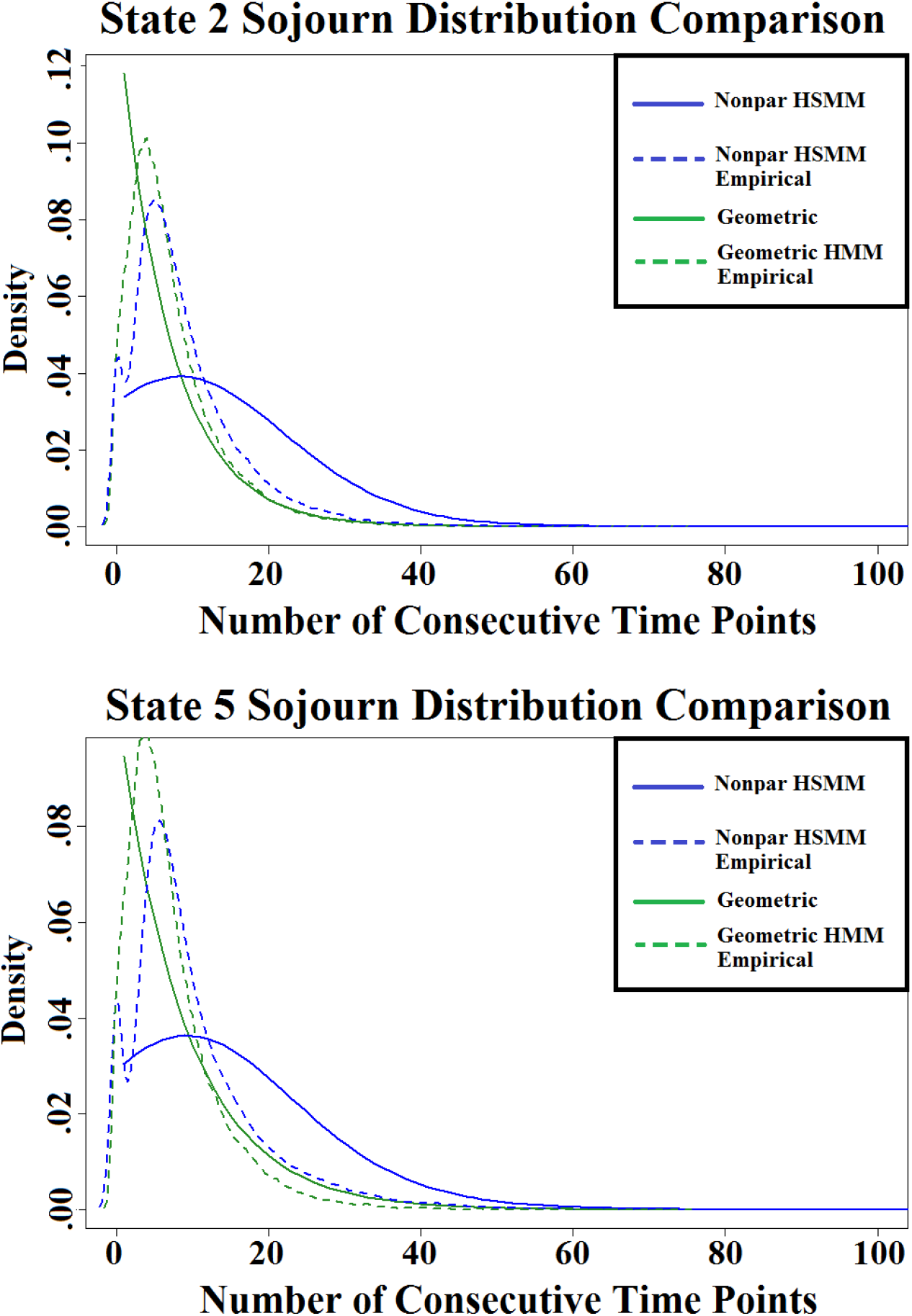
Comparison of sojourn distributions for two states under the standard HMM and the HSMM with a smoothed-nonparametric sojourn density. Plots include the direct sojourn distributions obtained from estimating the model parameters, as well as the empirical distributions calculated from estimating each subject’s most probable sequence of latent networks.

Table 1 reports the proportion of time spent in each state under all three models, as well as the average number of state switches and average number of consecutive time steps. As expected, the HMM produces a higher average number of state switches and smaller average number of consecutive steps compared to the HSMMs. The sojourn distributions for all 12 states that are estimated by each model choice are presented in the supplementary material.

**Table 1:** Proportion of time in each state under all three models, as well as the average number of switches and average number of consecutive time steps in a state.

|  | HMM | HSMM- Nonpar | HSMM- Pois |
| --- | --- | --- | --- |
| **State 1** | 0.086 | 0.086 | 0.099 |
| **State 2** | 0.113 | 0.120 | 0.137 |
| **State 3** | 0.105 | 0.100 | 0.022 |
| **State 4** | 0.067 | 0.070 | 0.075 |
| **State 5** | 0.120 | 0.118 | 0.112 |
| **State 6** | 0.109 | 0.105 | 0.111 |
| **State 7** | 0.060 | 0.061 | 0.080 |
| **State 8** | 0.046 | 0.056 | 0.054 |
| **State 9** | 0.065 | 0.057 | 0.071 |
| **State 10** | 0.068 | 0.070 | 0.072 |
| **State 11** | 0.086 | 0.081 | 0.092 |
| **State 12** | 0.075 | 0.075 | 0.075 |
| **Switches** | 139.441 (31.61) | 113.037 (25.90) | 130.010 (24.96) |
| **Consecutive Steps** | 9.406 (4.19) | 11.690 (5.64) | 9.738 (2.76) |

#### 3.3.4. Sojourn Distributions are Associated with Sustained Attention Scores

The participants in the HCP study also underwent a battery of behavioral tests. Here we focus on continuous sustained attention, as we anticipate that it may be related to transition time between states. It is measured using the Short Penn Continuous Performance Test (Number/Letter Version) [42, 43], in which participants view vertical and horizontal red lines flash on a screen. In one block, they must press a button when the lines form a number, and in the other block when the lines form a letter. The lines are displayed for 300 *ms* followed by a 700 *ms* inter-stimulus-interval. Each block contains a total of 90 stimuli and lasts for 1.5 minutes. The median response time for true positive responses was used as our outcome variable of interest.

We assess the relationship between sustained attention and the sojourn distribution of various brain states inferred from the HSMM assuming a non-parametric sojourn distribution. Our analysis was performed on a subset of 200 subjects from the HCP data set. These were chosen as the 100 subjects with the highest median response times on the task, as well as the 100 subjects with the lowest response times.

We first combined the 200 subjects and fit a HSMM to obtain 12 brain states common to both groups. We next modified the *MHSMM* R package in a way that would allow us to re-fit a HSMM to both groups separately while fixing the states to be those inferred from the combined analysis. In other words, the new HSMM fit to each group only re-estimated the transition matrix, initial state probabilities, and sojourn distributions (*u*_1:*K*_ and Σ_1:*K*_ remained the same).

After we obtained separate sojourn distributions for all 12 states, for each group, we performed a group comparison of each state’s sojourn distribution using a symmetrized version of the Kullback-Leibler (KL) divergence. KL divergence is a measure of the directed divergence between two probability distributions. The symmetrized version is defined to be

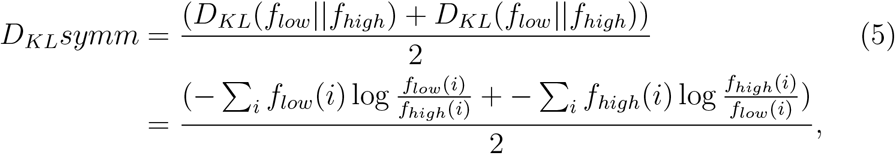

where *F_1ow_* and *F_high_* are the distribution functions of the low and high scoring groups respectively. In the simple case, a KL divergence of 0 indicates that we can expect very similar behavior from both distributions, whereas a KL divergence of 1 indicates that the two distributions behave in such a different manner that the expectation given the first distribution approaches zero. The *Philentropy* R package was used to calculate the KL divergence.

To determine if the difference we are seeing is meaningful, we performed a permutation test. We randomly permuted the group labels on the *n* = 200 subjects 500 times, re-fit the HSMM with the states fixed, re-calculated the KL divergence for each state and each of the 500 samples, and calculated the p-values for each state given this null distribution.

The estimated sojourn distributions for three different states (states 1, 3, and 12) are shown in Figure 10. These three states yield different results between groups, whereas the other states showed very minimal (if any) differences. In each of these 3 states, subjects with low sustained attention scores have a higher density for quicker switches out of the state. In other words, they do not sustain the state for as many consecutive time points compared to those with high scores. Figure 10 also shows the permutation test results for the three states. We see that the differences we are observing in sojourn distributions between the groups are unlikely to occur by chance (p-values 0.046, 0.032, and 0.026). Of course, adjusting for multiple testing, where the number of tests is 12, would yield larger p-values. However, we present these results as an illustration of the ability to extract potential meaningful information from the sojourn distribution.

**Figure 10:**
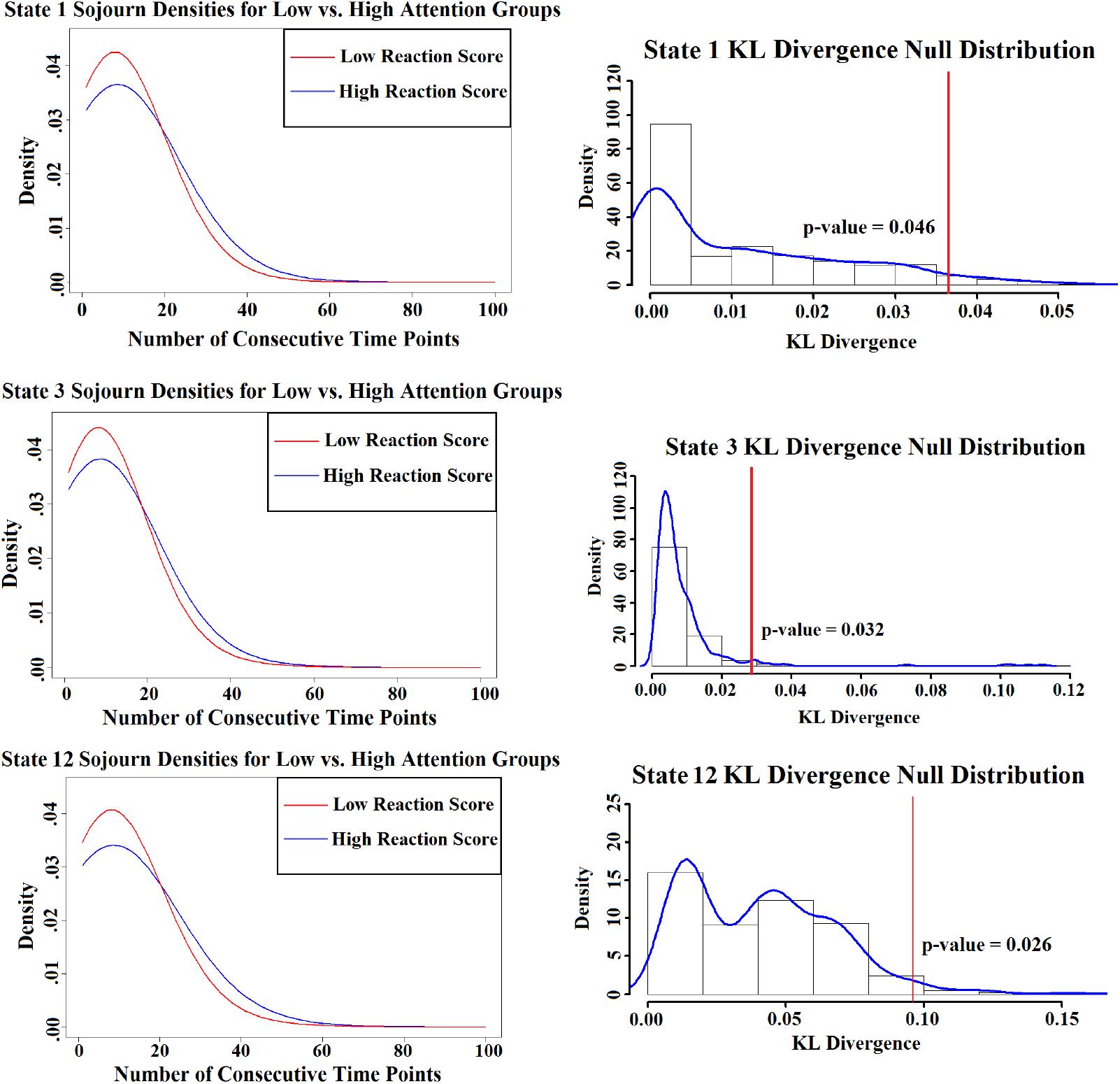
(Top row) Estimated sojourn distributions of the high and low sustained attention scoring groups for three states that revealed group differences. (Bottom row) Permutation test results obtained from permuting group labels and computing KL divergence between the estimated sojourn distributions for each group. The vertical line in each plot depicts where the observed group KL divergence group fell with respect to the null distribution.

Figure S8 in the supplement presents the three estimated states. States 1 and 3 are more similar, with the DMN regions appearing to be highly connected when looking at the Pearson correlation matrices. State 12 has many connections both within and between visual, somatomotor, and dorsal/ventral attention regions. The partial correlation matrices have more similarities between the 3 states, but state 12 appears to have slightly stronger partial correlations within the visual regions. The fact that the two groups of participants are yielding different sojourn distribution estimates on 3 of the 12 states seems to indicate that the sojourn distribution is an important aspect of dynamic FC analysis. For this reason, the HSMM has an advantage compared to the standard HMM given that the model directly estimates the sojourn distribution for each state without forcing the sojourn times to follow a potentially inaccurate geometric distribution.

Next, we explored the difference between the empirical sojourn distributions between the two attention groups. We took the same results used from fitting a HSMM on each group separately (with the 12 states fixed), but we estimated and compared the empirical sojourn distributions for each group, as opposed to the sojourn distributions estimated directly from the likelihood function and the E-M algorithm. The results of our permutation test (presented in Figure S9 in the supplementary material), show that the empirical sojourn distributions between the two groups differ for state 1 (p-value = 0.01). However, none of the other states’ sojourns distributions significantly differ between the two groups. These results support our prior conclusion that the attention groups differ in their sojourn times for state 1, given that they agree with our previous results. However, questions remain as to why we aren’t seeing empirical evidence of a difference in states 3 and 12. The increased noise from having to perform multiple steps to arrive at the empirical sojourn times (i.e. estimating the HSMM parameters, estimating the most probable sequence of states for each subject, and then estimating the empirical density from these state sequence estimates) may potentially be the reason, but this is something that will need to be explored in future work.

We also investigated whether there were any differences between groups in the geometric sojourn distributions fit directly from a standard HMM. We did not find any significant difference between the two groups in the estimated sojourn distributions nor did we see any group difference when comparing the empirical sojourn distributions for each state after fitting a standard HMM. We believe this is potentially due to the geometric distribution not being the best fit to the data, and therefore obscuring any real differences in sojourn times.

## 4. Discussion

The study of functional connectivity (FC), or the undirected association between two or more fMRI time series, has come to the forefront of research efforts in the field of neuroimaging. While many of the brain connectivity analyses performed to date estimate a single static network structure for the length of time a subject is scanned, there is growing evidence that valuable information is obtained by estimating several networks over shorter time periods. In other words, it is desirable to estimate at which time points subjects are switching network states, as well as the network states, themselves.

There are several existing approaches towards assessing time-varying connectivity. These include using sliding window correlations [6], change point models [8, 9, 10], wavelet transform coherence [5], time series models [11, 12], and switching vector autoregressive models [13, 15]. In recent years, hidden Markov models (HMMs) have also proven to be a useful modeling approach towards assessing FC. HMMs make the assumption that the time series data at each brain region can be characterized via a series of hidden/latent brain states. However, HMMs are limited by their implicit assumption that the sojourn distribution, the distribution of the number of consecutive time points in a state, is geometrically distributed. When this assumption is not an accurate representation of the data, one may obtain an inaccurate estimate of the state sequence on a subject-by-subject basis. More specifically, the model will place an unrealistic amount of weight on quicker state switches.

In this paper, we propose a more principled and flexible approach using hidden semi-Markov models (HSMMs) when the true distribution of the sojourn times is either unknown or suspected to not be geometrically distributed. That is, we propose using HSMMs to find each subject’s most probable series of network states and the graphs associated with each state. The HSMM does not implicitly assume a geometric sojourn distribution for each state. Rather, it allows one to explicitly model and estimate the sojourn distributions by inserting an additional term into the likelihood function.

In some task-based experimental paradigms, it may be the case that a geometrically distributed sojourn distribution is well motivated and is a reasonable fit for the data. For example, one could imagine a task requiring subjects to rapidly switch attention, where it would be beneficial to assume a higher probability for quick transitions. However, in many experiments, this may not be the case. In the anxiety provoking experiment [27] described in this paper, we found the Poisson distribution to be a reasonable model for the data. It may also be the case that one does not have a good intuition to what the sojourn distribution may look like, such as in the resting-state fMRI case. Therefore, in these instances, as long as the amount of data is sufficient for the number of parameters, we recommend working with a non-parametric sojourn distribution. In cases where the sample size is not very large, it is advisable to only use the nonparametric sojourn distribution as a rough estimate. One could get an idea of a parametric distribution to use based off of this estimate and then fit the model with that distribution.

The impact of an incorrectly specified sojourn distribution, such as with the standard HMM, can potentially be far reaching. What we have chosen to focus on in this paper is the difference in estimated sojourn times and state switches when fitting models with various sojourn distributions. We illustrate that if one is interested in accurately estimating subject-level state switches, correctly modeling the sojourn distribution is important. We have also shown the importance of obtaining accurate estimations of the sojourn distributions if one wishes to associate them with group behaviors and traits. However, we have not extensively explored how an incorrectly specified sojourn distribution may impact other aspects of the model fit. Our simulation study suggests a misspecified sojourn distribution may not impact the estimation of states, but may impact the estimate of transition probabilities between states to some extent. However, the seriousness of this impact in various settings will need to be further explored in future work.

We believe our model can be improved upon by incorporating random effects. A limitation is that we are fitting one set of model parameters for all subjects. However, it would be useful to have subject-specific states, sojourn distributions, and state transition probability matrices. In this setting, estimates would be calculated in a manner that borrows strength from the group, but also allows for individual differences. We plan to develop a mixed-effects HSMM framework in future work.

To conclude, the HSMM framework affords a flexibility that the HMM framework does not. It allows for direct modeling and estimation of the sojourn distribution for each state instead of assuming a geometric distribution, which has the potential to significantly impact results when estimating the state sequence on a subject-by-subject basis. We demonstrate that the HSMM approach gives promising results, as indicated by the near perfect capturing of the anxiety inducing experimental design, as well as accurate estimation of model parameters in our simulation study. We make comparisons between the standard HMM approach and the HSMM approach throughout the paper, illustrating that results *do* differ depending on the model used. The flexibility of the HSMM compared to the standard HMM in obtaining accurate estimates of sojourn times has the potential to play an important role in understanding healthy and diseased brain mechanisms.

## 5. Acknowledgements

This work was supported by NIH grants P41 EB015909 and R01EB016061, as well as the Johns Hopkins University Provost’s Postdoctoral Fellowship Program.

