## Supplement for "Improved state change estimation in dynamic functional connectivity using hidden semi-Markov models"

*Compiled January 12, 2019*

**The study of functional brain networks has grown rapidly over the past decade. While most functional connectivity (FC) analyses estimate one static network structure for the entire length of the functional magnetic resonance imaging (fMRI) time series, recently there has been increased interest in studying time-varying changes in FC. Hidden Markov models (HMMs) have proven to be a useful modeling approach for discovering repeating graphs of interacting brain regions (brain states). However, a limitation lies in HMMs assuming that the sojourn time, the number of consecutive time points in a state, is geometrically distributed. This may encourage inaccurate estimation of the time spent in a state before switching to another state. We propose a hidden semi-Markov model (HSMM) approach for inferring time-varying brain networks from fMRI data, which explicitly models the sojourn distribution. Specifically, we propose using HSMMs to find each subject's most probable series of network states and the graphs associated with each state, while properly estimating and modeling the sojourn distribution for each state. We perform a simulation study, as well as an analysis on both task-based fMRI data from an anxiety-inducing experiment and resting-state fMRI data from the Human Connectome Project. Our results demonstrate the importance of model choice when estimating sojourn times and reveal their potential for understanding healthy and diseased brain mechanisms.**

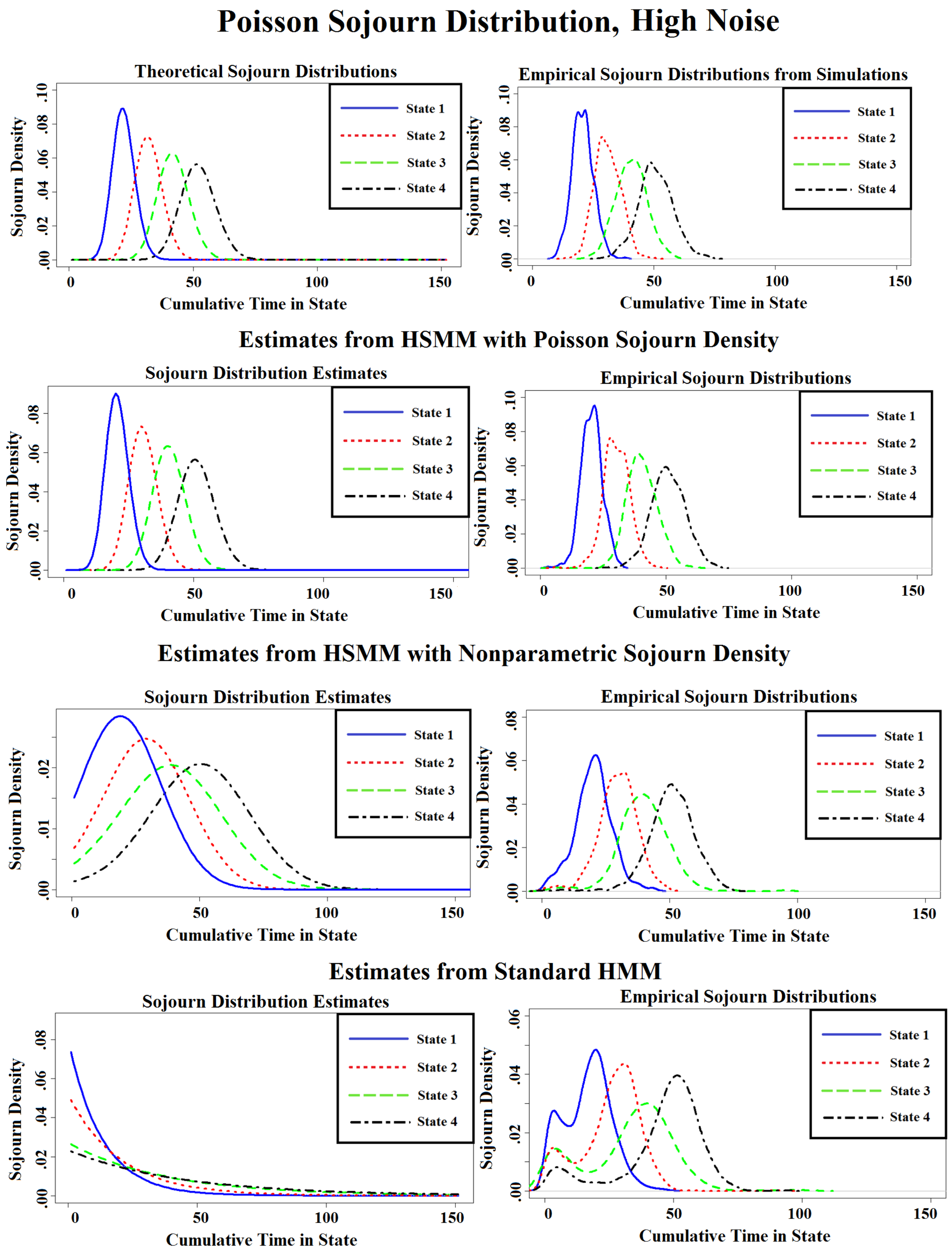


**Fig. S1.** Sojourn distributions, as well as empirical sojourn distributions, that were estimated via the HMM and HSMMs under the simulation scenario of Poisson sojourn distributions and high noise.

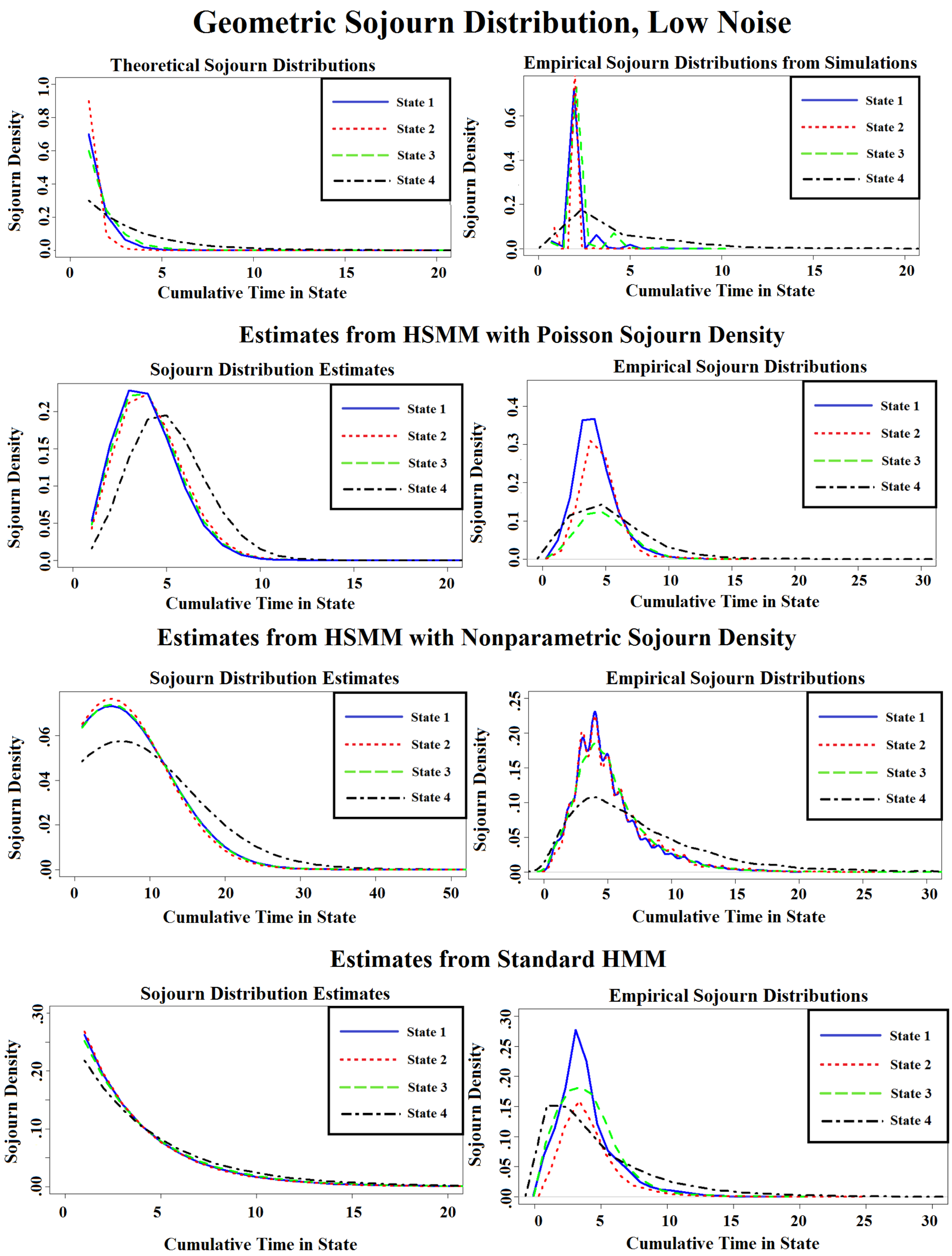


**Fig. S2.** Sojourn distributions, as well as empirical sojourn distributions, that were estimated via the HMM and HSMMs under the simulation scenario of Geometric sojourn distributions and low noise.

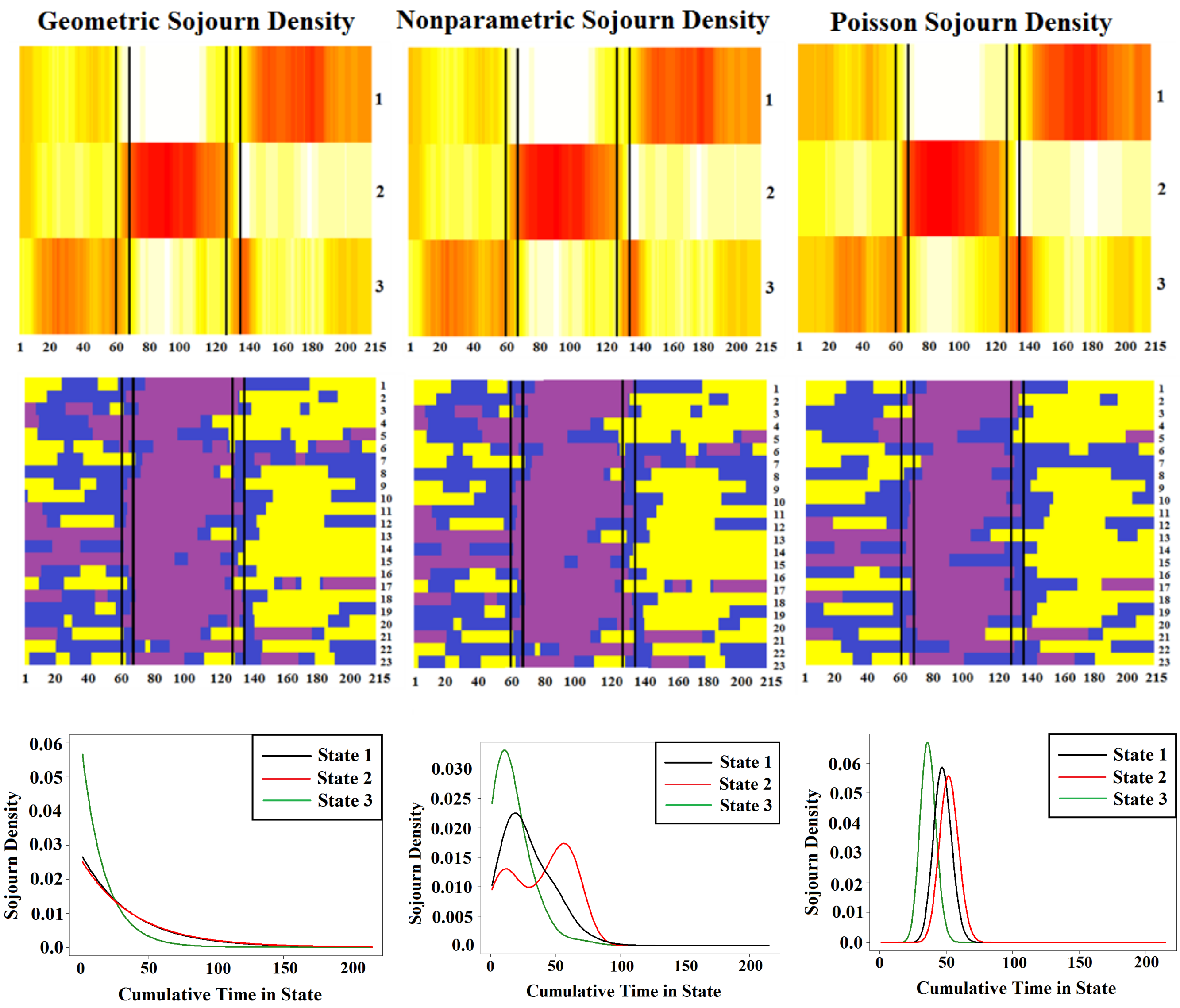


**Fig. S3.** (Top row) Heatmaps representing number of subjects in each state across scan time. (Middle row) Estimated state sequence for each subject. State 1: yellow, State 2: purple, State 3: blue. (Bottom row) Estimated state sojourn distributions for each model.

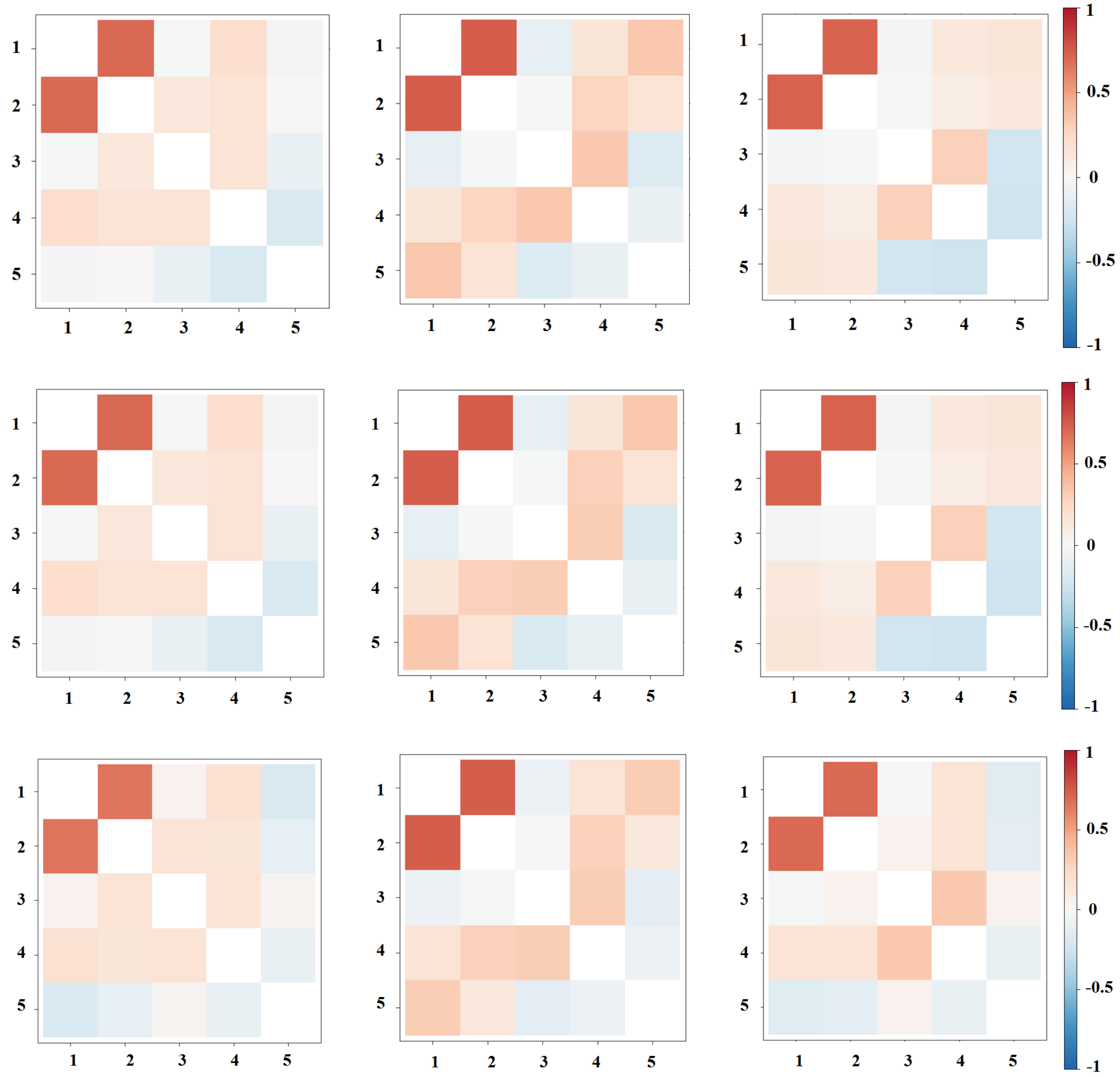


**Fig. S4.** Pearson correlation matrices representing the three network states (States 1, 2, and 3 respectively) for each model. Nodes (1-2) represent two different subsections of the ventral medial prefrontal cortex (VMPFC), (3) the ventral striatum, (4) the dorsal lateral prefrontal cortex (DLPFC); and (5) heart rate. Dark red indicates a large positive correlation, while dark blue indicates a large negative correlation.

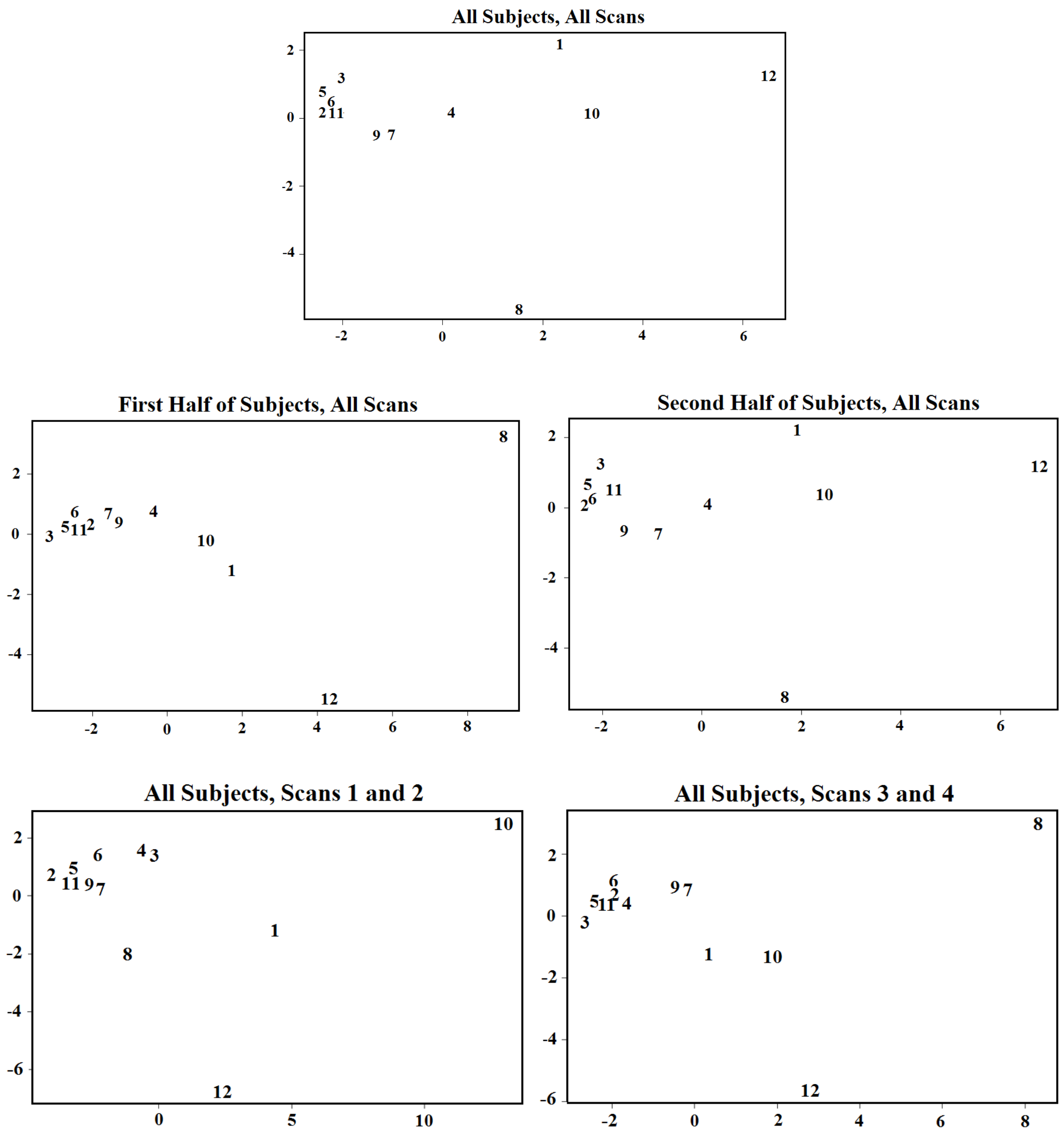


**Fig. S5.** Multidimensional scaling representation of the 12 network states. Relative position of one state compared to another state is determined by the Euclidean distance between covariance matrices. (Top) Scaling is done on networks found for the HSMM fit on all subjects and across all scans. (Middle) Scaling is done on networks found for the HSMM fit on the first and second half of the subjects separately. (Bottom) Scaling is done on networks found for the HSMM fit on the first two scans and last two scans separately.

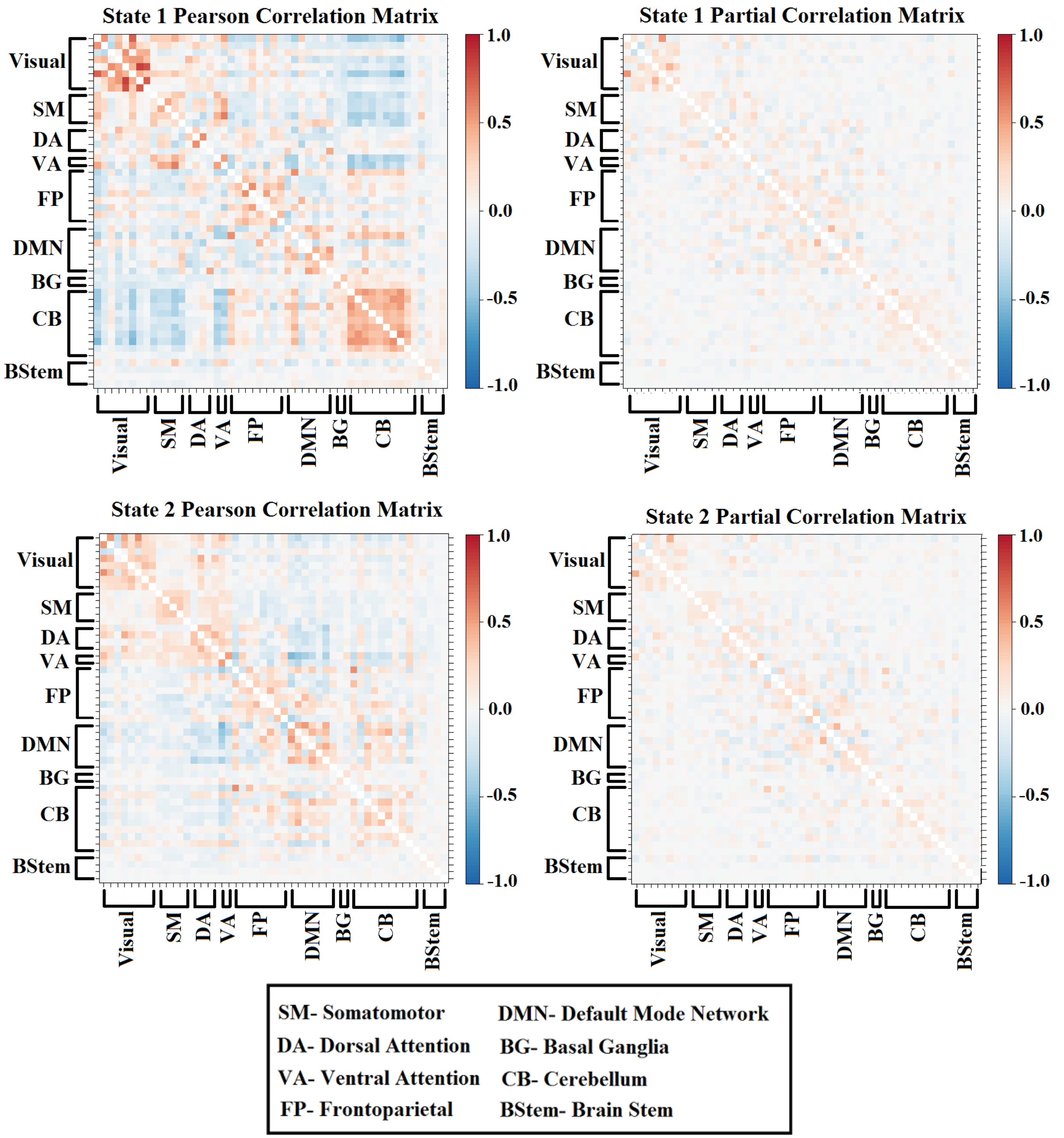


**Fig. S6.** Pearson and partial correlation matrices representing states 1-2 for the HSMM with the smoothed-nonparametric sojourn distribution.

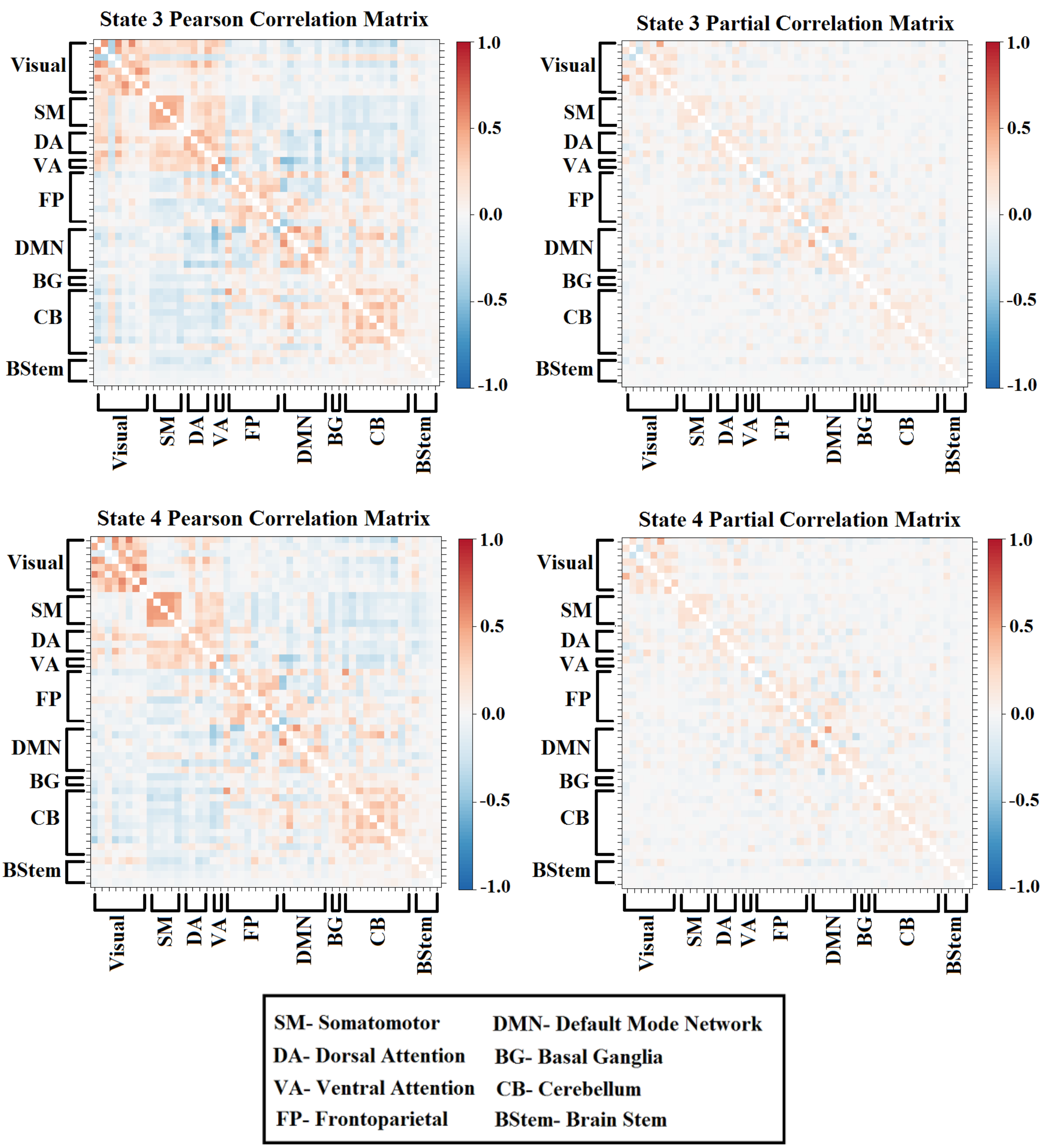


**Fig. S6.** Pearson and partial correlation matrices representing states 3-4.

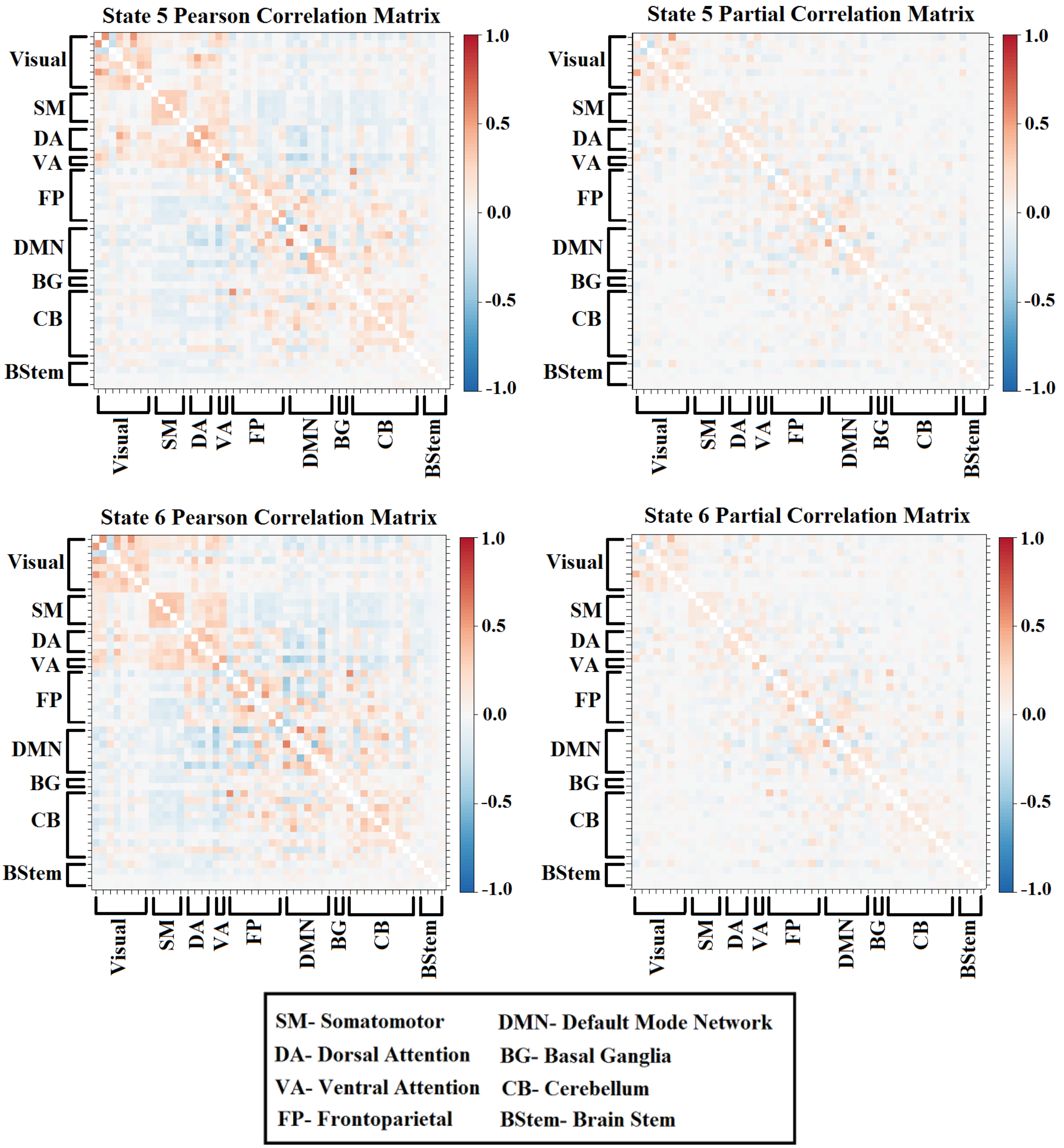


**Fig. S6.** Pearson and partial correlation matrices representing states 5-6.

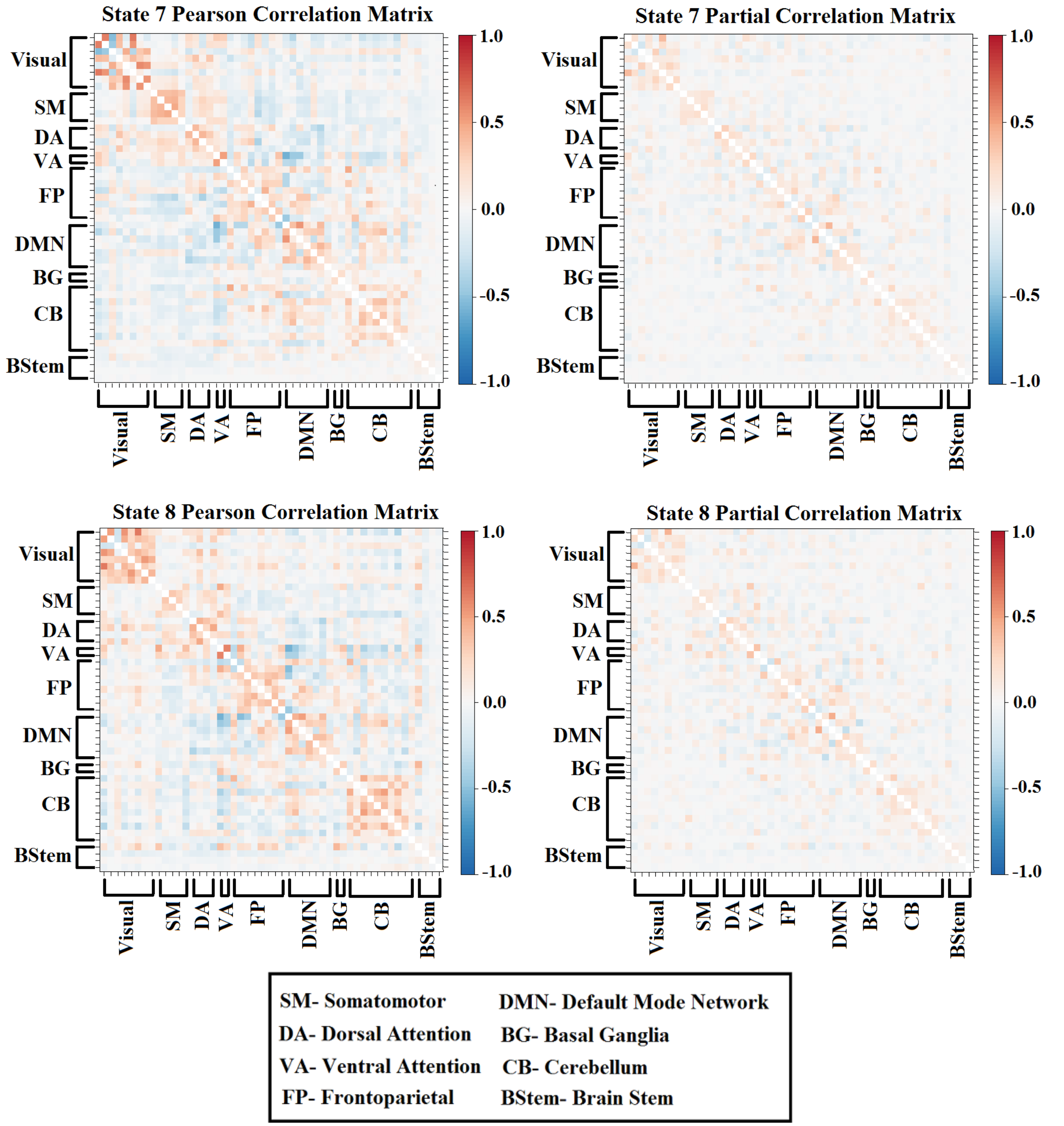


**Fig. S6.** Pearson and partial correlation matrices representing states 7-8.

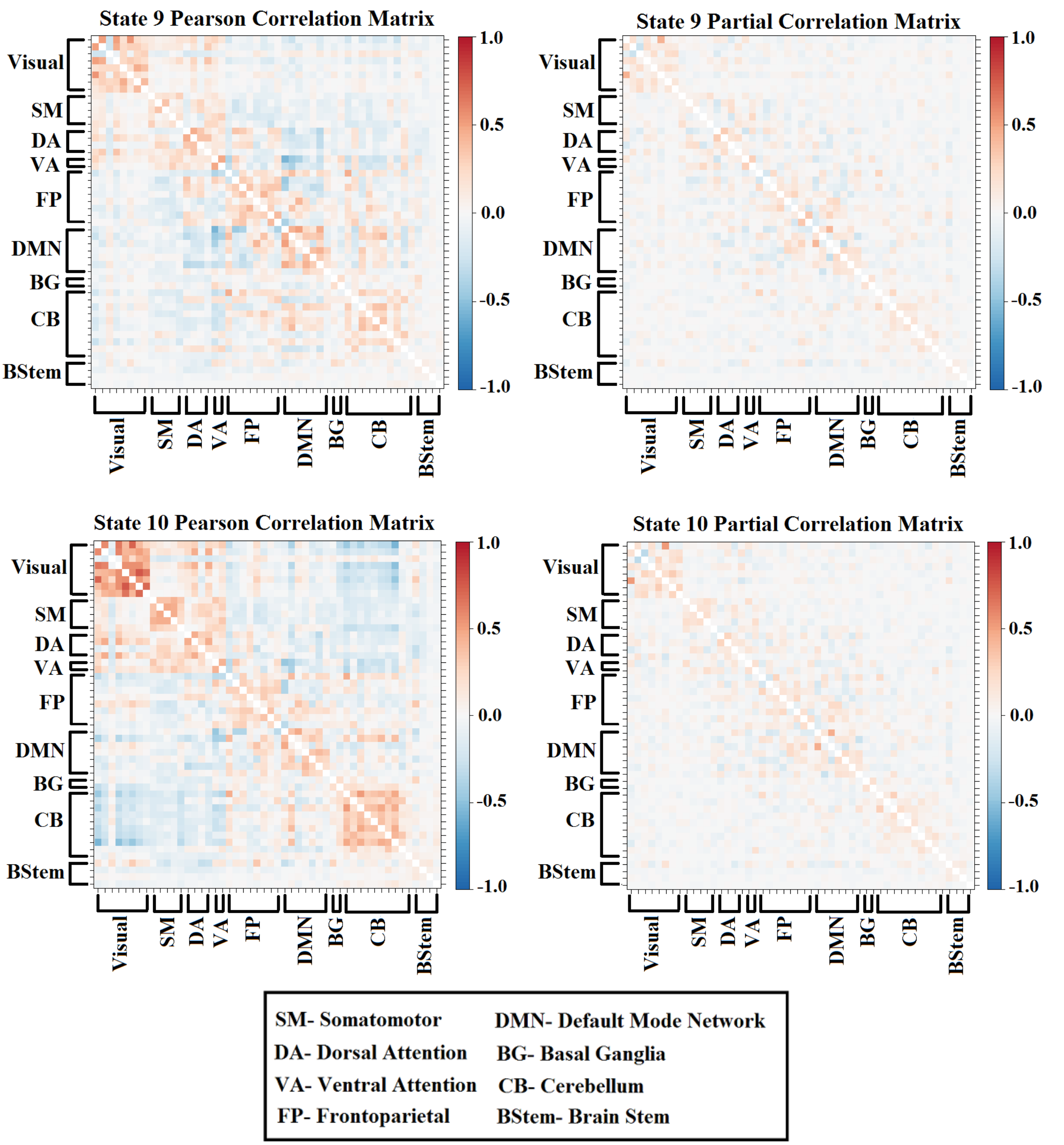


**Fig. S6.** Pearson and partial correlation matrices representing states 9-10.

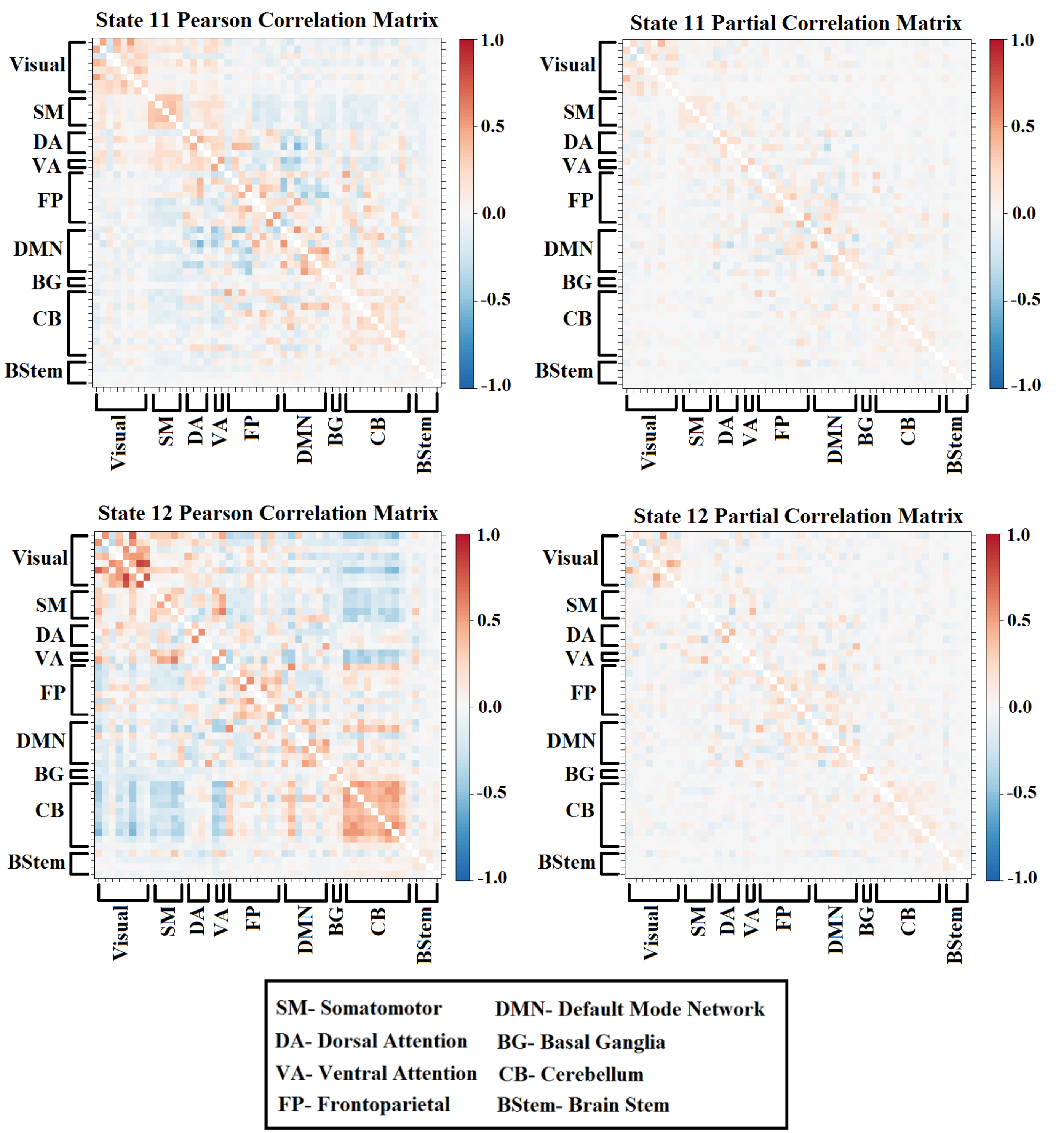


**Fig. S6.** Pearson and partial correlation matrices representing states 11-12.

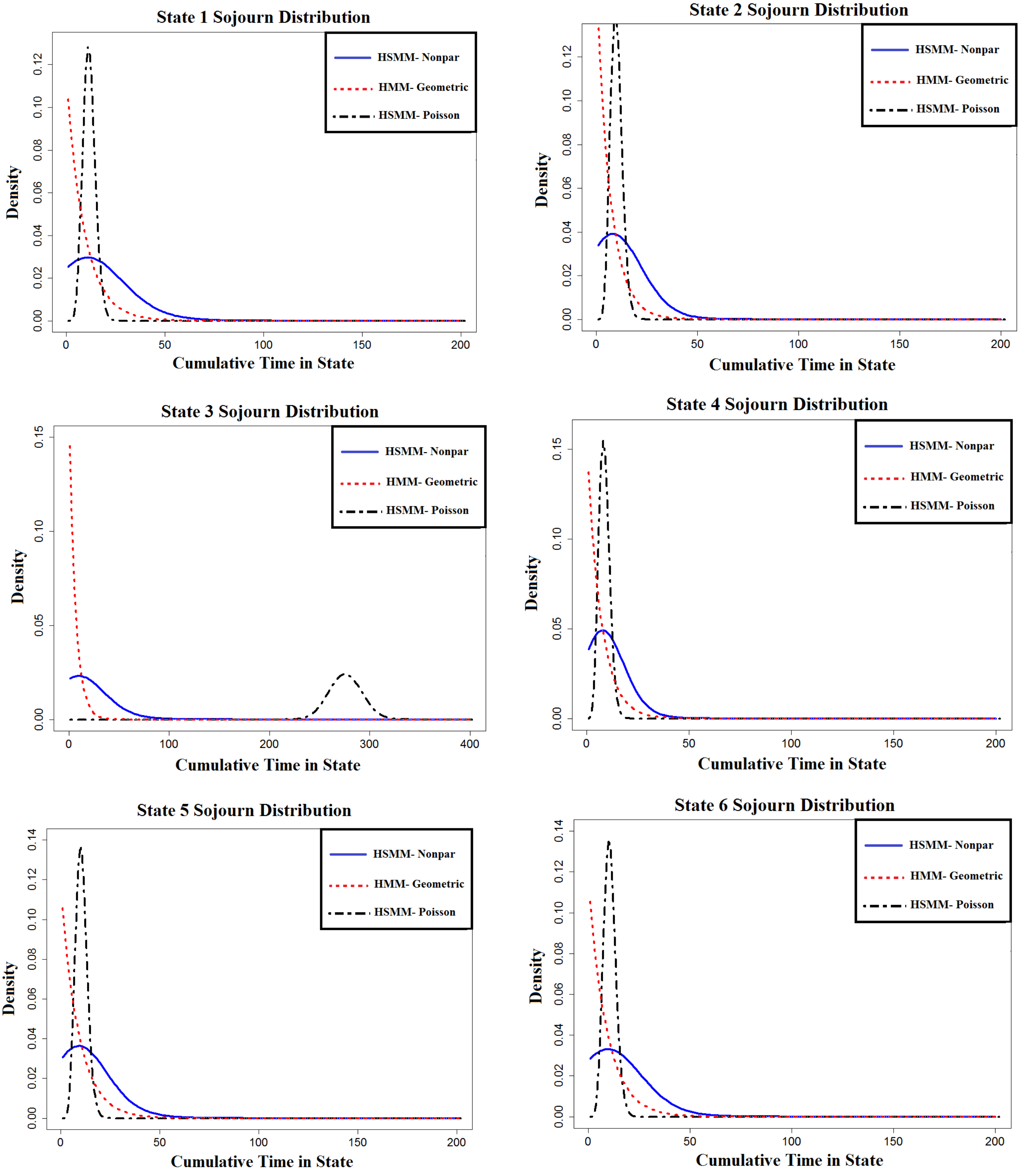


**Fig. S7.** Sojourn distributions estimated from the HCP data under the two HSMMs and the standard HMM for states 1-6.

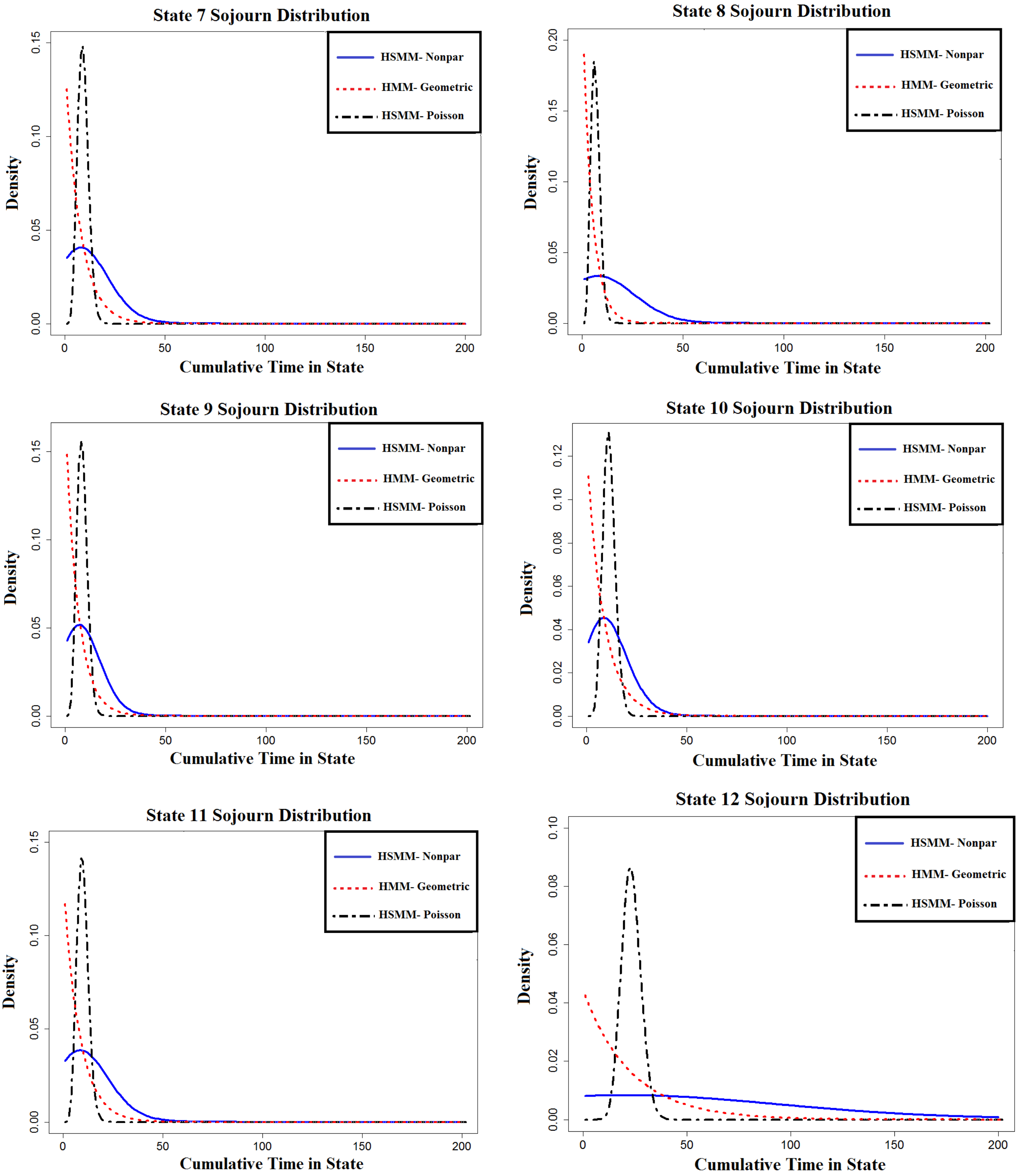


**Fig. S7.** Sojourn distributions estimated from the HCP data under the two HSMMs and the standard HMM for states 7-12.

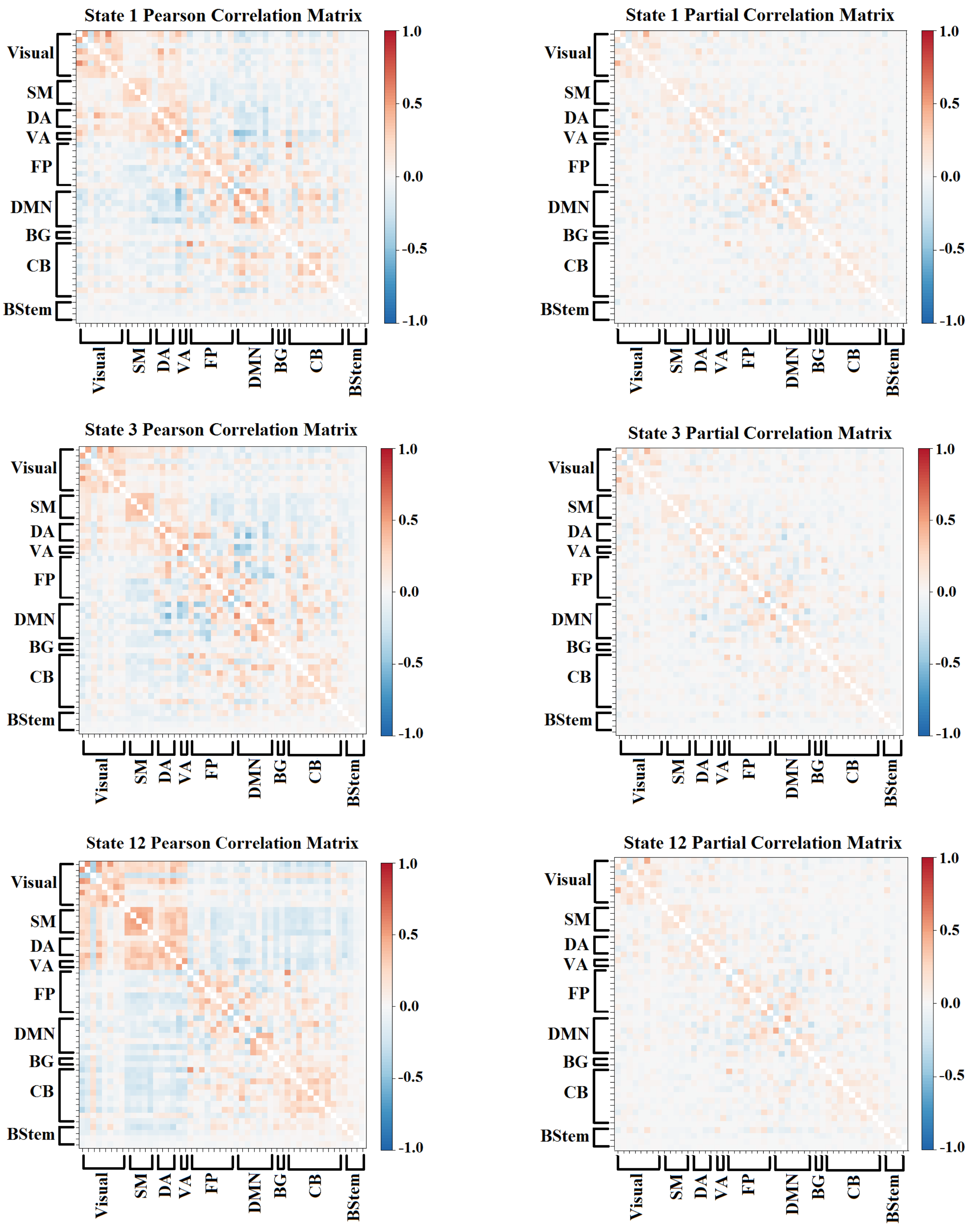


**Fig. S8.** Pearson (left column) and partial (right column) correlation matrices for the three states with sojourn distributions that differ between the high and low sustained attention scoring groups. Abbreviations are as follows: SM- Somatomotor, DA- Dorsal Attention, VA- Ventral Attention, FP- Frontoparietal, DMN- Default Mode Network, BG- Basal Ganglia, CB- Cerebellum, BStem- Brain Stem.

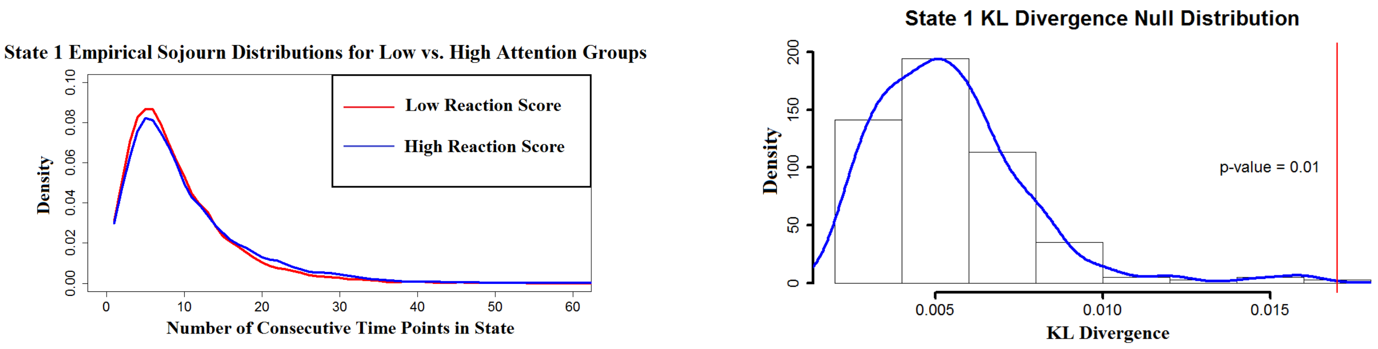


**Fig. S9.** (Left) Estimated empirical sojourn distributions of the high and low sustained attention scoring groups for the one state that revealed group differences. (Right) Permutation test results obtained from permuting group labels (200 permutations) and computing KL divergence between the estimated sojourn distributions for each group. The vertical line depicts where the observed group KL divergence group fell with respect to the null distribution
